# Competitive Adsorption of a Monoclonal Antibody and Amphiphilic Polymers to the Air-Water Interface

**DOI:** 10.1101/2024.09.27.615546

**Authors:** Elise J. Hingst, Michaela Blech, Dariush Hinderberger, Patrick Garidel, Christian Schwieger

## Abstract

Understanding structure and self-organization of monoclonal antibodies (mAbs) at the air-water interface is important for the stability and effectiveness of protein drug formulations used in pharmaceutical industry. This paper investigates the competitive adsorption of a mAb and the amphiphilic surfactants poloxamer 188 (P188) and polysorbate 20 (PS20), both of which are commonly used to prevent mAb surface adsorption. Firstly, it is studied whether these surfactants prevent mAb adsorption, and secondly, whether it is possible to desorb mAb molecules from the air-water interface by surfactant addition. For surface pressure and surface tension data, Langmuir film balance measurements and drop shape tensiometry were used. Infrared Reflection-Absorption Spectroscopy (IRRAS) provided information on the surface composition, including the amount of adsorbed molecules. P188 exists in different self-assembled phases depending on its surface concentration. Our experiments show that the phase state of P188 has a significant impact on mAb adsorption. The presence of P188 in the brush phase (≥ 0.3 mg/L) consistently inhibits mAb adsorption. On the contrary, addition of P188 after mAb film formation could not cause desorption of mAb. However, addition of PS20 leads to desorption of freshly formed interfacial mAb layers of up to two hours age. Interestingly, an aged mAb layer of 17 hours could not be desorbed by PS20. This suggests a time dependent reorganization of mAb at the air-water interface, which increases its resistance to desorption. These findings are discussed with respect to possible inter-molecular interactions within the interfacial film.

## INTRODUCTION

Monoclonal antibody (mAb) solutions, particularly at high concentrations, play an important role in pharmaceutical industry.^1-6^ Subcutaneous injection has become a common way to administer immunoglobulins ^7^ and offers a number of advantages compared to intravenous injection, including economy of time,^8^ cost saving,^8^ and the possibility for simple and independent home therapy.^9^ The structure of the extracellular matrix limits the subcutaneous injection volume to less than 2 mL.^10^ Therefore, either several subcutaneous injections or a highly concentrated product is required ^7^. MAb concentrations used in pharmaceuticals are up to 300 mg/mL.^10^

Due to their, inter alia, physicochemical properties, formulations of such highly concentrated mAb solutions are very challenging.^2^ On the one hand, there is a tendency towards protein particle formation,^11^ which can occur at almost every step during mAb production, e.g., manufacturing or storage.^12^ Several factors like buffer conditions, temperature, light and presence of surfaces may foster protein aggregation.^2^ On the other hand, the high viscosities of highly concentrated mAb solutions hamper drug formulation development. Hence, syringeability often becomes problematic for subcutaneous injection.^1, 13^ Furthermore, higher treatment time and injection pain may occur.^14, 15^ Changes in product stability could lead to reduced efficacy and unwanted immunogenic responses, including anaphylaxis.^13^ To impede these reactions and processes and improve formulations, understanding drug stability and mAb behaviour in concentrated solution is of high relevance.^5^ Previous studies show the influence of electroviscous effects,^16, 17^ pH,^11, 16^ buffer conditions,^18, 19^ and added excipients like hydrophobic^18^ and polar salts,^19^ mono- and disaccharides,^11^ and amino acids^20^ on highly concentrated mAb solution viscosity and stability.

Another important factor is the presence of several interfaces.^21^ MAb adsorption to SiO_2_/water interfaces,^22^ which are a good model for glass/water interfaces, as well as adsorption to in-line filters used for intravenous application^23^ were observed. Especially, interactions of mAb with the air-water interface are the topic of current research.^24, 25^ In particular the effect of spontaneous mAb adsorption on the efficacy of the protein drug formulation is of high interest. Protein particle formation and unfolding processes at the air-water interface are not fully understood^21^ and it cannot be ruled out that the formulation stability and safety is affected by these processes. Hence, strategies to generally impede mAb adsorption to the air-water interface are required. Previous studies have shown that surfactants used as additives or excipients are successful in preventing protein self-assembly at the air-water interface.^21^ Poloxamer 188 (P188) and polysorbate 20 (Tween®20, PS20) are two of the U.S. Food and Drug Administration (FDA) approved surface-active excipients.^26^ Polysorbates (PS) can be used for drugs administered by the main injection routes (subcutaneous, intravenous as well as intramuscular) and represent the major surfactant added to protein drug formulations.^3, 27^ Various studies aimed at revealing the mechanisms behind the stabilizing effect exerted by these excipients in highly concentrated mAb solutions.^4, 6, 25^ Two main paths have been suggested: (i) intermolecular interaction of surfactant molecules with exposed hydrophobic mAb-sites (direct binding) and (ii) surfactant adsorption at the air-water interface with consequential surface shielding for mAb adsorption.^3, 28^ Experiments show that mAb stability due to addition of PS is predominantly caused by PS blocking the surface^6, 11^ and not by bulk interactions.^4^ Due to possible enzymatic degradation processes^29^ and its effect on fatty acid aggregation,^30^ there is need for other amphiphilic excipients as alternatives to PS.^31^ So far, P188 represents an alternative surfactant for protein drug formulations.^27^

In this paper, we present results from competitive air-water interface adsorption studies involving PS20 and P188 in the presence of mAb. Adsorption kinetics were studied using Langmuir film balance measurements and drop shape tensiometry. Infrared reflection-absorption spectroscopy (IRRAS), was used to derive information on the molecular composition of the mAb/surfactant interfacial films at various surfactant concentrations and film formations procedures.^28^ We conclude on conditions where the mAb adsorption to the air-water interface can be prevented or already formed mAb films can be replaced from the interface.

## MATERIALS AND METHODS

### Materials

The structures of the used substances are shown in Scheme 1. ChemDraw 22.2.0 and PyMOL 2.3.3 were used as graphic software. The monoclonal antibody (mAb) used in this study was provided by Boehringer Ingelheim Pharma GmbH & Co. KG (Biberach, Germany). It belongs to the class of IgG1 antibodies. Its structure is similar to but not identical with the structure shown in Scheme 1A. The average molar mass of the used mAb is 146’000 g/mol. Poloxamer 188 (P188, Scheme 1B) and polysorbate 20 (PS20, high purity qualities, Scheme 1C) were obtained from BASF and Croda International, respectively. P188 has an average molecular weight of 8’500 g/mol. The average molecular weight of PS20 is 1228 g/mol. Chloroform, ethanol and methanol (all HP grade) were obtained from Carl Roth GmbH & Co.KG (Karlsruhe, Germany). L-Histidine and L-Histidine · HCl · H2O were obtained from Ajinomoto OmniChem (N.V., USA). Ultra-pure Milli-Q water (κ < 0.055 μS/cm) was used for all preparations and experiments.

**Scheme 1.**
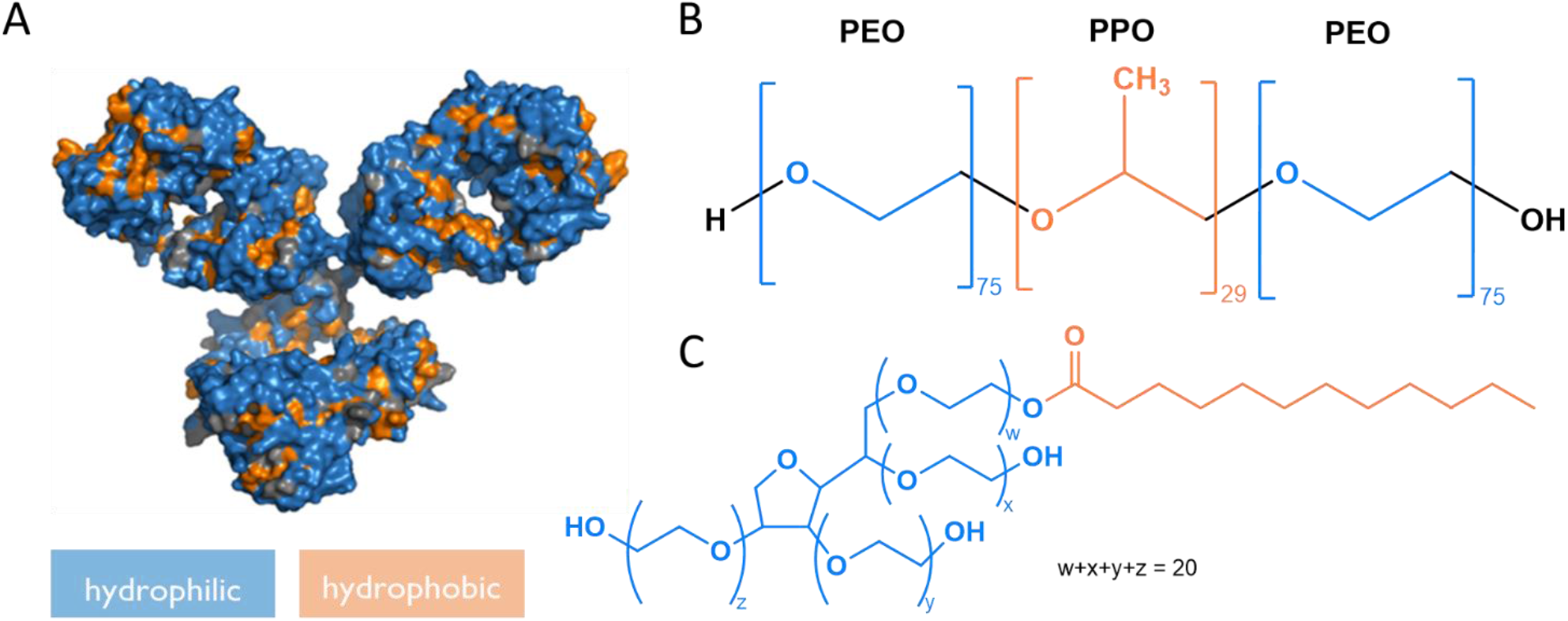
(Chemical) structures of tested substances with color-coded hydrophilic (blue) and hydrophobic (orange) moieties. A: An example of an antibody structure (pdb: 1IGY), B: The triblock copolymer P188, PEO: Poly(ethylenoxide), PPO: Poly(propylenoxide). C: Structure of poly(ethylenoxide) sorbitan lauric acid ester, PS20.

### Sample Preparation

For preparation of protein and surfactant solution, the required amount of components were weighed using an analytical balance and subsequently dissolved in 24.4 mM histidine buffer (11.4 mM L-Histidine, 12,8 mM L-Histidine · HCl · H_2_O, pH = 6.0 ± 0.2). This buffer composition is used for all preparations and experiments described in this paper.

P188 solutions were dissolved in a concentration range from 0.5 mg/L (approximately 0.06 μM) to 20 g/L (approximately 2.4 mM) in histidine buffer and mixed on a vortex mixer. For the measurement of compression isotherms, a 0.122 mM solution was prepared by dissolving P188 in chloroform.

For preparation of PS20 solutions, PS20 was dissolved in a concentration range from 1.2 mg/L (approximately 0.001 mM) to 12.28 g/L (approximately 10 mM) in histidine buffer and mixed by gentle agitation due to strong foaming.

Mab solutions were diluted in a concentration range from 50 mg/L (approximately 0.3 μM) to 20 g/L (approximately 0.1 mM) in histidine buffer and mixed by gentle agitation. The stock solution had a concentration of 90 g/L.

### Monolayer Preparation

Distinct volumes of proteins and surfactants dissolved in 24.4 mM histidine buffer (pH = 6.0 ± 0.2) were carefully injected into the histidine buffer subphase of the thoroughly cleaned and filled film balance trough using glass syringes (Hamilton Bonaduz, Bonaduz, Switzerland). All measurements were performed at 20 °C.

### Langmuir film balance measurements at the air-water interface

All Langmuir film balance measurements were performed on two home-built circular PTFE-troughs (Scheme S1) with a volume of about 10 mL. A perspex sheath protects the surface from draught and dust. Four cellulose- and water-filled bowls under the sheath ensured a constant protein concentration in the subphase by securing constant air humidity and preventing subphase-evaporation. The surface pressure was measured by a microbalance (Riegler und Kierstein GmbH, Berlin). A filter paper (Wilhelmy plate) was used as surface pressure probe. All measurements were performed at 20 °C ± 0.1 °C, regulated by a water bath (Thermostat F6, Haake, Karlsruhe, Germany).

Before and after each measurement the troughs were cleaned with Hellmanex™ (Hellma GmbH & Co. KG, Müllheim) in a 1:100 dilution for at least 20 minutes exposure time and subsequently with ultra-pure water. A fresh Wilhelmy plate was attached after every experiment involving protein. A clean trough is characterized by a rupture of the water layer without droplet formation when aspirating the solution.

The surface tension σ of ultra-pure water (72.8 mN ·m^-1^) ^32^ and the surface tension of air (0 mN ·m^-1^) at 20 °C were used as reference for calibrating the pressure sensor. First, water was filled into the trough and the device-specific fill level was adjusted using a height-standardised cannula. Adjusting the specific fill levels (± 0.5 mm) for each device is necessary to obtain optimal, reproducible results. Subsequently, the surface pressure π was regulated to the value of 0 mN ·m^-1^. Calibration at the second point was performed by suctioning the subphase until the Wilhelmy plate was no longer covered, where π was set to of 72.8 mN ·m^-1^. Note that the surface pressure is calculated as 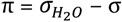.

Swift operation is important as otherwise, the calibration value would be distorted by evaporation effects. Once the calibration was completed, histidine buffer was filled into the trough, and the temperature was allowed to stabilize before adjusting the device-specific filling height again. After calibration, the troughs filled with histidine buffer were equilibrated for 20 minutes.

The sample solution was injected through a lateral hole in the trough or through a lockable aperture in the sheath, depending on the injected component. PS20 was injected through the sheath to prevent trough-leakage and consequential adulteration of the fill level. During the measurement, the subphase was gently stirred by a magnetic stirrer.

### Infrared Reflection-Absorption Spectroscopy (IRRAS) at the air-water interface

A scheme of the instrument is shown in the Supporting Information (Scheme S2). IRRAS measurements were performed on a BRUKER Vector 70 FT-IR spectrometer including an A511 reflection unit (Bruker Optics, Ettlingen, Germany) and a liquid nitrogen cooled MCT detector. The Langmuir trough setup (Riegler & Kierstein, Potsdam, Germany) contains a circular sample trough with a diameter of 6 cm, equipped with a Wilhelmy plate as pressure probe, and a reference trough (30 × 6 cm^2^). In both troughs the temperature was held constant at 20 °C by a thermostat (F6, Haake, Karlsruhe, Germany). Using a shuttle-system either trough can be brought to the focus of the IR beam as needed. Defined and constant filling levels in the troughs are important to achieve reproducible results. This is ensured by a computer controlled pumping unit connected to a laser reflection signal which provides the filling level data. To enable a stable atmosphere a perspex sheath was placed over the Langmuir trough setup and the IR-reflection unit.^33^

Both troughs were filled with the same buffer subphase and the filling level was adjusted as described above. Prior to sample injection, up to ten buffer-buffer spectra were recorded to check the purity of the trough and the surface. After that, 5 μL of the sample solution were gently injected into the subphase at two different positions and in the lane of the magnetic stirrer.

The angle of incidence of the IR-beam can be varied between 25 ° and 70 °. The polarization of the IR-beam can be varied between perpendicular (s) or parallel (p), with respect to the plane of incidence. Time dependent series of spectra were obtained in steps of 5-minute-intervals between two sample measurements. All adsorption measurements were conducted with an s-polarized IR-beam, an incidence angle of 40 °, a scan number of 1000 scans per spectrum, a resolution of 8 cm^-1^ and a zero-filling factor of 4. The penetration depth of the IR-beam is 0.5 – 1.0 μm. For recording and initial analysis the spectroscopy software OPUS (Bruker Optics) was used. Isotherms were analysed and plotted with the software OriginPro 2019.

### Spectral Simulations and Fits

First, an offset was subtracted from the measured spectra such that the wavelength region between 2500 cm^-1^ and 2600 cm^-1^ was set to an average of zero. This was followed by subtraction of averaged spectra of the pure buffer to compensate for atmospheric influences. For best compensation of atmospheric and buffer influences, the averaged buffer spectrum was scaled such that after subtraction the variance of the second derivative was minimal. Since the H_2_O bending vibration (1650 cm^-1^) partially overlaps with the amide I band at 1658 cm^-1^, a computer simulated spectrum of a non-adsorbing layer with a thickness of 1 nm and pure water as a subphase with its optical parameter taken from Bertie *et al*. was subtracted.^34^ The simulated spectrum was scaled such that after subtraction the total intensities were minimal in the range of 3680 cm^-1^ to 3450 cm^-1^. From the scaling of the simulated spectrum, we were able to estimate absolute values of the layer thickness of adsorbed layers in nm. Eventually, characteristic bands of the compounds present at the interface (Table S1) were integrated to obtain a measure for their interfacial concentration.

### Drop Shape Tensiometry

The drop profile analysis tensiometer PAT-1M by SINTERFACE Technologies (Potsdam, Germany) in the pendant drop mode was used for adsorption experiments. Prior to every experiment, tubes were rinsed with ultra-pure water. To examine adequate purity, the surface tension of a water drop (V = 35 μL) was measured. The surface tension must stay constant in a range of 71.0 – 73.0 mN ·m^-1^ for at least a 5 m minutes. If this was not the case, the system was rinsed with a 1 % Hellmanex™ solution and with ultra-pure water again. Remaining water was rinsed off the tube and the sample was aspirated. The first two to three drops of the sample solution were discarded. All measurements were performed with a cannula of 3 mm diameter and a drop volume that was held constant at 20 μL. After finishing the experiment, the system was rinsed with ultra-pure water. The software PAT-1M was used. Data points were saved every second, which includes values of surface tension (mN ·m^-1^), drop area (mm^2^), drop volume (mm^3^) and profile error. For calibrating the device, camera calibration was performed using a spherical stainless-steel probe (d = 3 mm). The focus can be adjusted in the setup of the program as well as with a setting wheel at the device.

## RESULTS AND DISCUSSION

### Surface activity of pure compounds

Before addressing the surfactant-mAb interactions at the air-water interface, the pure components were measured separately for proper referencing. We used a combination of drop shape tensiometry and Langmuir film balance measurements to study of the surface activities of the components as a function of their bulk concentration. Both methods are complementary and are used to obtain cross-validated results.

Drop shape tensiometry (pendant drop) needs much lower sample volumes (about 200 μL) than Langmuir film balance measurements (up to 10 mL). Furthermore, drop shape tensiometry enables experiments with high bulk concentrations (> 20 g/L) and long-time measurements extending to several days (with adaptive drop volume or drop area control via computer and camera). Both types of experiments cannot be realized with the film balance. On the contrary, film balance measurements are more sensitive and can be used for very low concentrations (< 1 mg/L). Since very small changes in surface tension resulting from low volume concentrations do not lead to detectable variations of the drop shape, drop shape tensiometry is not suitable for measurements below 1 mg/L.

### P188 behavior at the air-water interface

P188 exists in different self-assembled phases at surfaces, depending on its surface concentration, ^35, 36^ which are shown in Scheme 2. Injecting P188 molecules into water leads to adsorption of the molecules to the air-water interface due to the hydrophobic PPO-blocks (orange coloured in Scheme 2). Since the inherently hydrophilic PEO blocks (blue coloured) also possess a certain degree of hydrophobicity, they also accumulate at the air-water interface at low surface concentrations, forming flat structures, referred to as “pancakes”.^35^ Increasing surface concentration reduces the available space, causing the EO units to be displaced from the surface into the subphase. Thereupon, mushroom shaped structures (“mushroom” phase) form, followed by brush-like structures (“brush” phase). These phase states have been described previously in general for amphiphilic block copolymers by Haefele *et al*..^37^ Furthermore, Haefele *et al*. describe a film organization state called “cigar”, which follows the “brush” state. With our methods we were not able to detect a higher condensed state than “brush”, because no further plateau above π = 11 mN ·m^-1^ is clearly discernible in the compression isotherm (see Figure 1). Consequently, we refer to the highly condensed phase as the “brush” phase hereinafter. A compression isotherm of P188 is shown in Figure 1, which relates to previous experiments.^36, 38^ The “mushroom”-”brush” transition can be observed in the compression isotherm as a pseudo-plateau at 11 mN/m, whereas the “pancake”-”mushroom” transition is gradual. Herein, we refer to P188 films between 5 and 11 mN ·m^-1^ as P188 “mushroom” phase.

**Scheme 2.**
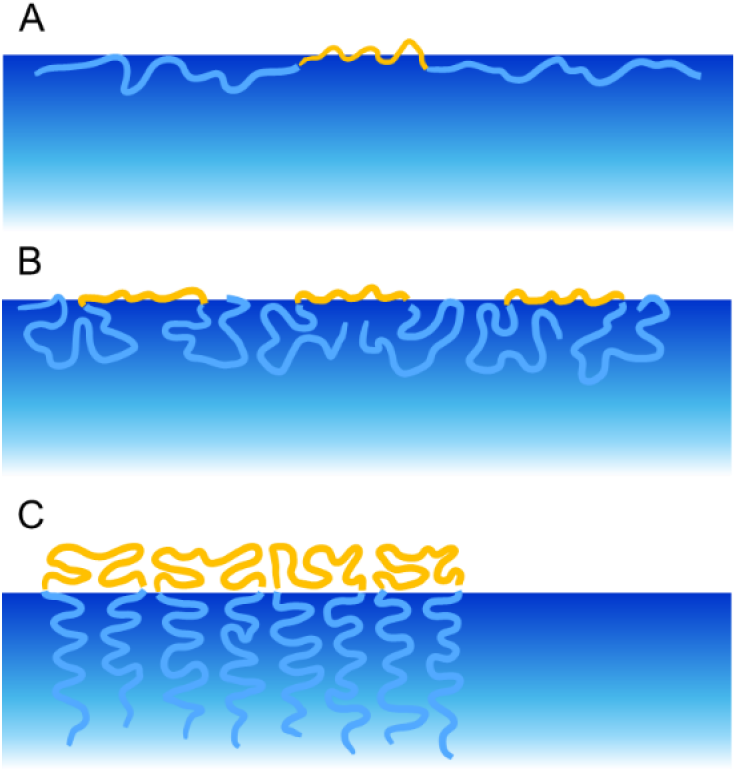
Schematic representation of the behaviour of P 188 at the air-water interface with increasing surface pressure adapted from ^37^ . At low surface pressures, the surfactants lie flat on the surface, while at higher pressures, the PEO blocks protrude into the bulk phase. A: “pancake” phase, B: “mushroom” phase, C: “brush” phase. The hydrophobic PPO blocks are orange-coloured, while the hydrophilic PEO blocks are blue-coloured

**Figure 1.**
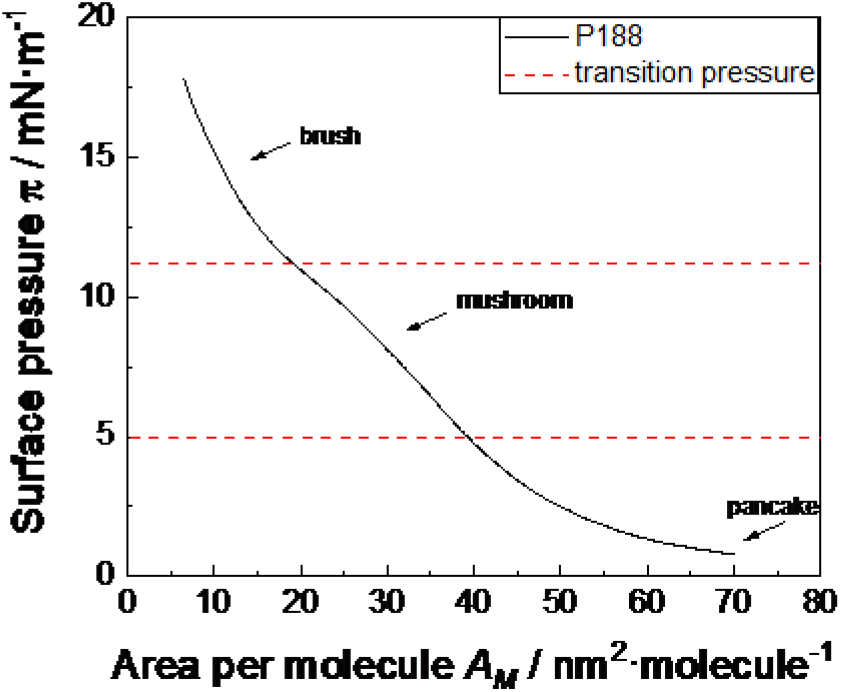
Compression isotherm of P188, performed on histidine buffer (pH = 6.0 ± 0.2) at a temperature of T = 20 °C. The dotted red lines show the transition pressure from “pancake” to “mushroom” at π = 5 mN ·m^-1^ and from “mushroom” to “brush” at π = 11 mN ·m^-1^.

### Adsorption behavior

The adsorption isotherms (Figure 2) show quite different adsorption kinetics for the three studied components when they are injected into the aqueous subphase. Both P188 and PS20 are more surface active and adsorb much faster to the interface than the mAb: within only 2 minutes, a stable surface pressure π is attained. P188 reaches the saturation equilibrium pressure at 18 mN ·m^-1^ ^38^ and PS20 at 35 mN ·m^-1^ ^28^, indicating a higher surface activity of PS20 when compared to P188.

**Figure 2.**
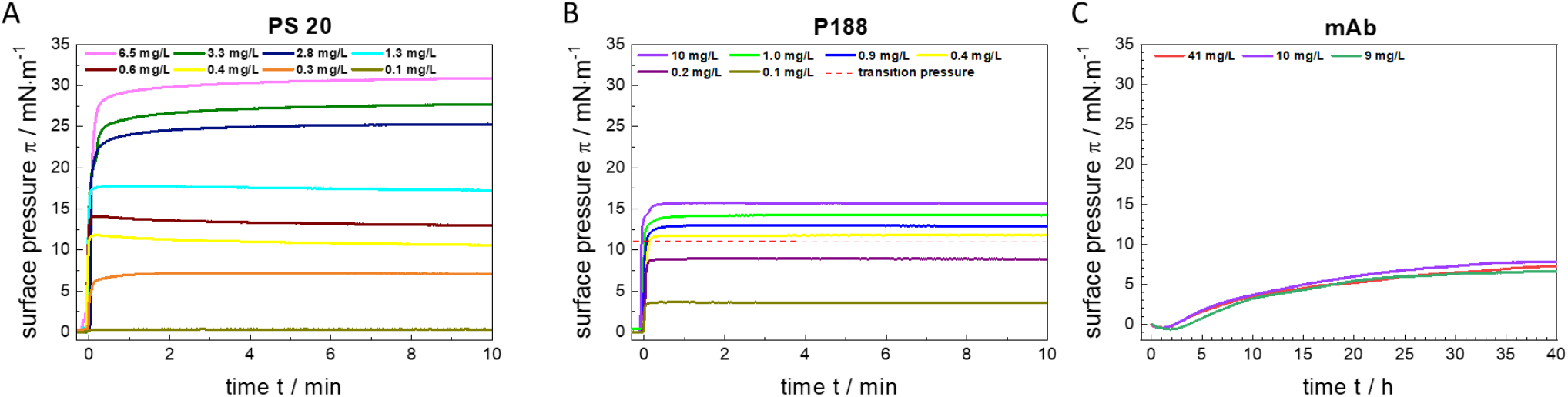
Adsorption isotherms of polysorbate 20 (PS20), poloxamer P188 and a monoclonal antibody (mAb) at different subphase concentrations determined by Langmuir film balance method. Measurements were performed on histidine buffer (pH = 6.0 ± 0.2) at a temperature of T = 20 °C. Note the identical surface pressure axes scales while the time axes are different. B: The upper red dashed line shows the transition pressure of the “mushroom”-phase to the “brush”-phase at π = 11 mN ·m^-1^, obtained from the compression isotherm in Figure 1. The lower red dashed line shows the transition pressure of the “pancake”-phase to the “mushroom”-phase at about π = 5 mN ·m^-1^.

The surface behaviour of the mAb is shown in Figure 2 C. The difference in adsorption kinetics compared to the surfactants is striking. MAb adsorption proceeds slower, i.e., the equilibration takes several hours and the mAb surface activity (ca. 7 mN/m) is clearly lower than that of the surfactants. This can be explained by the much lower amphiphilic character of the mAb. While P188 and PS20 have clearly discernible hydrophilic and hydrophobic moieties, hydrophobic areas of the mAb are distributed throughout the large protein molecule and partially hidden inside the structure. These hidden areas may be exposed at the surface in case of adsorption which would be one reason for the significant differences in kinetics and activity as compared to the surfactants.^39, 40^ The slow adsorption kinetic of the mAb can in addition be explained by its high molecular weight of about 146’000 g/mol. Compared to P188 (8’500 g/mol) and PS20 (1’228 g/mol), the mAb is a large molecule which diffuses much slower to the air-water interface. By molecular dynamic simulations a radius of gyration of 4.1 ± 0.87 nm per P188 molecule was obtained.^41^ The radius of gyration of a PS20 micelle was determined to be 2.1 – 2.2 nm by molecular dynamic simulations ^42^ which agrees well with experimental results obtained by dynamic light scattering (2.4 nm).^43^ MAb gyration radii values are at about 5 nm, calculated by Guinier analysis.^44^

In addition, we performed drop shape tensiometry measurements at similar conditions (Figure S1). The results corroborate the findings obtained by the film balance measurements described above.

According to the generated data, clearly showing faster adsorption kinetics and higher surface activity of the investigated surfactants, these should be able to shield the surface and prevent mAb adsorption, either when being injected prior to or when injected simultaneously with the mAb. This is, however, only true when no synergistic effects and no interactions between mAb and the surfactants are assumed. In the following paragraphs the competitive adsorption of mAb and P188 as well as mAb and PS20 is investigated and discussed in detail.

### Competitive adsorption of monoclonal antibody and surfactants to the air-water interface

Competitive surface adsorption experiments were performed to study how mAb behaves in presence of either P188 or PS20. To this end, injection sequence, time of injection and concentrations were varied. The experiments were conducted on an adsorption Langmuir film balance. Furthermore, infrared reflection-absorption spectroscopy (IRRAS) was performed to obtain information on the molecular surface composition, which is possible since the different absorbing molecules lead to distinguished bands in the absorption-reflection spectra.

All experiments were performed with a sub-phase composed of 24.4 mM histidine buffer (pH = 6.0 ± 0.2). The experiments were conducted with concentrations below the critical micelle concentration (*cmc*) of 0.48 mM for P188 ^45, 46^ and 0.059 mM for PS20^47^. Plots for calculating the cmc can be found in the Supplementary Information (Figure S2).

In the following we describe three different sets of experiments as schematically shown in Scheme 3. That is i) simultaneous injection of either surfactant and the mAb into the subphase ii) injection of mAb underneath a pre-formed film of surfactant, and iii) injection of the surfactant underneath a pre-equilibrated film of mAb. Thereby case i) simulats the situation where a surfactant und mAb are both present in solution and a new surface is created by shaking or handling a mAb formulation.

**Scheme 3.**
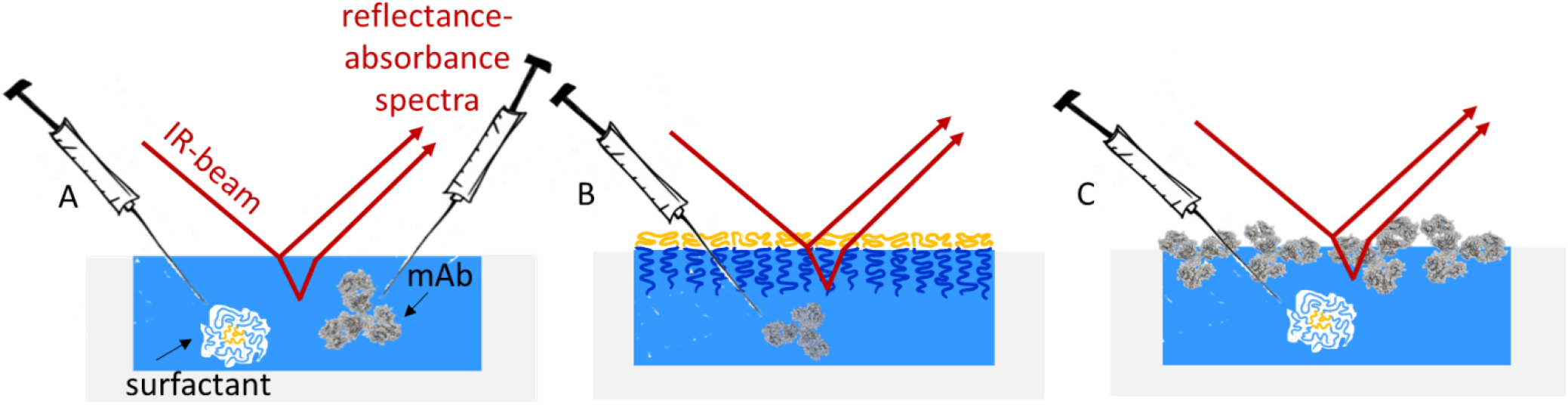
Used experimental setups. Surfactant constitutes either P188 or PS20. A: Simultaneous injection of surfactant and mAb into the pure buffer (24.4 mM histidine, pH = 6.0 ± 0.2), B: Injection of mAb underneath a preformed surfactant layer, C: Injection of surfactant underneath a pre-formed mAb layer.

In Figure 3, IRRA spectra of the pure components at the air-water interface are shown for reference. For each compound, unique characteristic reflection-absorption bands, that do not overlap with absorptions of the other compounds were chosen as reporter bands. Their integrals are used as measures of the amount of adsorbed component at the air-water interface. For PS20 that is the carbonyl stretching vibration ν(C=O), for P188 the asymmetrical ether stretching vibration ν(C-O-C) and for the mAb the amide I and amide II band. The water stretching vibration ν(HOH) originates from absorption of the aqueous subphase and is used for layer thickness determination as described in Schwieger et al. 2012. Before further evaluation, the subphase water contribution is eliminated from the spectra by subtraction of simulated IRRA spectra of a non-absorbing layer with adjusted layer thickness. The resulting water-compensated spectra are shown in Figure S3.

**Figure 3.**
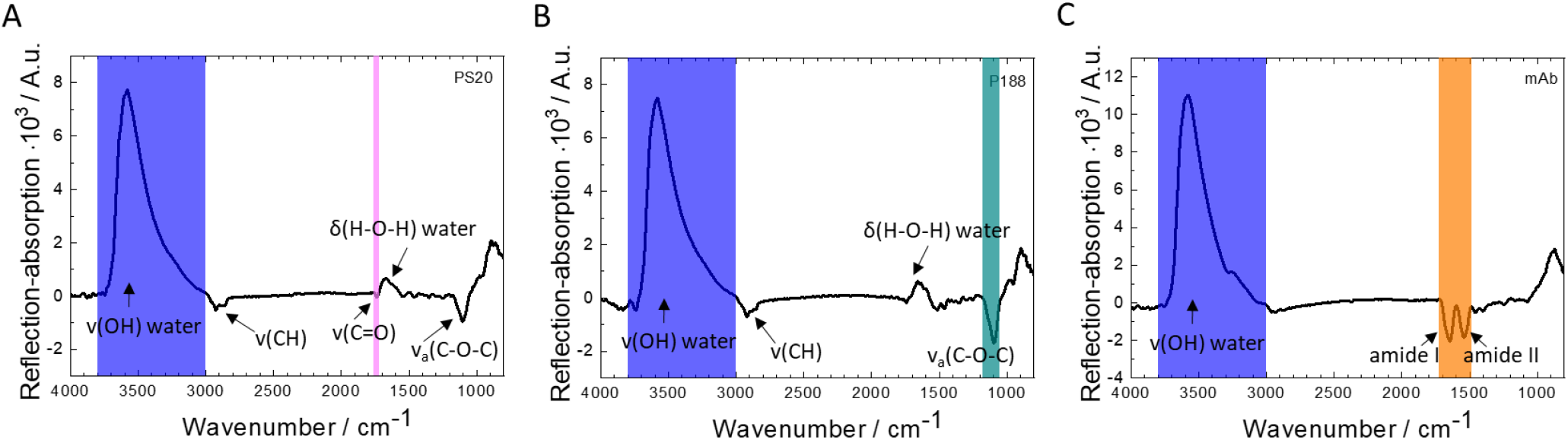
Infrared reflection-absorption (IRRA) spectra of the pure compounds at the air-water interface. A: PS20 after 30 minutes of adsorption, subphase concentration 1 mg/L, B: PS20 after 30 minutes of adsorption, subphase concentration 1 mg/L, C: after 15 hours of adsorption, subphase concentration 5 mg/L. ν, stretching vibration; δ, deformation vibration. Integration limits of the respective bands are shown as coloured, transparent boxes.

The here studied concentrations of P188, PS20 and mAb are well below those used in pharmaceutical biologics (18fold lower for mAb). However, the chosen molar ratios of surfactant to mAb coincide with those used in pharmaceuticals. Due to area requirement calculations for mAb (SI), we estimate that the air-water interface is almost entirely covered by mAb molecules in the here chosen experimental conditions. Therefore, higher bulk concentrations are not expected to lead to higher surface concentrations or different interactions with the surfactants at the interface. The here described adsorption behaviors and interactions between mAb and surfactants at the air-water interface are assumed to be representative for higher concentrated solutions.

#### 1. Simultaneous injection of surfactants and antibody

To study the direct competitive behavior of the components, IRRA spectroscopic measurements with simultaneous injection of both components, mAb and either P188 or PS20 at final bulk ratios of ≥ 1 or 2.6 surfactant molecules per antibody molecule, were performed. A mAb subphase concentration of 5 mg/L was used for each measurement. IRRA Spectra and time dependent integral intensities of the marker band for the experiments with P188 and mAb are shown in Figure 4. In this experimental setup a P188-phase dependent behavior becomes apparent.

**Figure 4:**
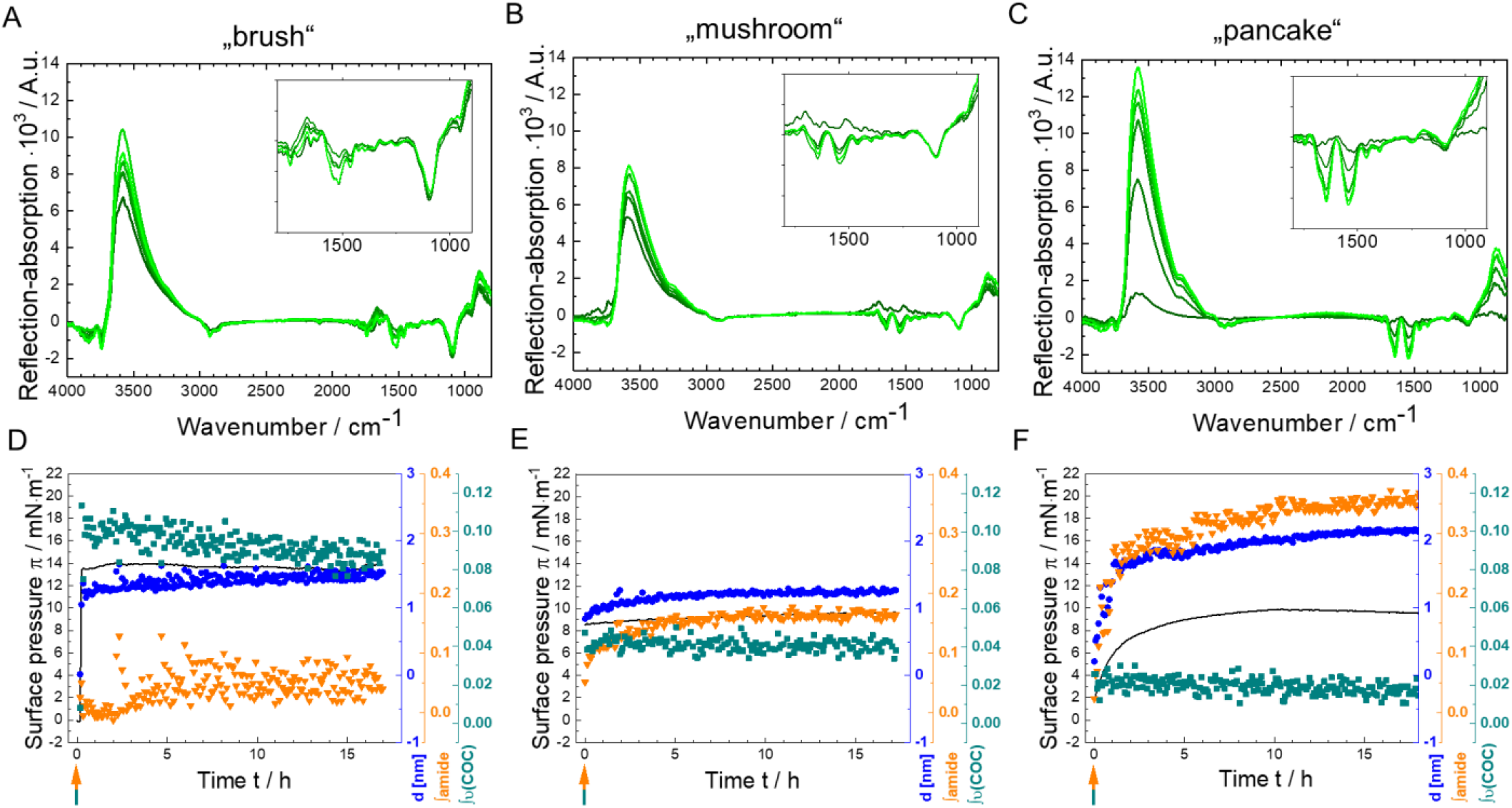
Competitive surface activity experiments P188 and mAb, using simultaneous Infrared reflection-absorption spectroscopy (IRRAS) and surface tension measurements. A simultaneous injection of mAb (*c*_sub_ = 5 mg/L) and P188 in different concentrations (A,D: “brush” 1 mg/L, B,E: “mushroom” 0.13 mg/L, C,F: “pancake” 0.11 mg/L) was performed. Measurements were performed on 24.4 mM histidine buffer (pH = 6.0 ± 0.2) at a temperature of T = 20 °C. A-C: IRRA spectra 5 minutes to 15.5 hours after mAb and P188 injection. The insets show an enlarged part of the spectra in a wavenumber range of 1800 cm^-1^ to 900 cm^-1^. The colour representation corresponds to different adsorption times. Darker shades of green indicate earlier spectra, while lighter shades of green indicate later spectra. D-F: Surface pressure π (mN ·m^-1^) (black line), layer thickness d 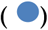, integral of amide I and amide II bands 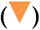 and integral of asymmetrical C-O-C-stretching vibrational band centred at 1100 cm^-1^ 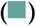 as a function of time (h). The green-orange arrow shows the injection time of P188 and mAb into the subphase.

When working with bulk concentrations that leads to P188 self-assembly in the “brush”-phase, only P188 adsorbs to the air-water interface and mAb is not able to co-adsorb, which can be seen in Figure 4A and D. Figure 4A shows the IRRA spectra of the measured P188-mAb-mix. Figure 4D shows the time dependent evolvement of surface pressure π (black line), layer thickness *d* (blue dots), integral of amide-I and amide-II-bands of the mAb (orange triangles) and the integral of the asymmetrical C-O-C-stretching vibration of P188 centred at 1100 cm^-1^ (green squares). The surface pressure increases immediately after the simultaneous injecting of P188 and mAb, reaches its maximum and remains constant over the time of the measurement. The fast increase in surface pressure as well as its equilibrium value of ca 15 mN/m is particularly attributable to the adsorption of P188, as evidenced by the results obtained from the pure component measurements described above. The constant surface pressure indicates a fast, unfluctuating surface layer formation. The average layer thickness is determined from the ν(HOH) vibration of the subphase water to be ∼1.5 nm and fits reasonably well with previous studies.^48^ Based on the amide and ν(C-O-C) band integrals, the average surface layer composition can be estimated. The ν(C-O-C) integral increases together with the surface pressure immediately with the injection to ∼0.1 mRA/cm, indicating the fast and stable adsorption of P188 to the interface. On the contrary, the amide band integral is zero or close to zero throughout the course of the experiment, indicating that the mAb cannot adsorb to the interface in these conditions. Correspondingly, in the associated IRRA-spectra (Figure 4A) the characteristic ν(C-O-C) vibration of P188 at 1100 cm^-1^ is strongly pronounced, whereas in the wavenumber range of the amide I and II bands at 1725 cm^-1^ to 1485 cm^-1^, only the deformation vibration of water δ(HOH) at 1650 cm^-1^ is discernible.

In the presence of P188 in concentrations leading to formation of the “mushroom”-phase, the time dependent integrals of the amide band and the ν(C-O-C) band (Figure 4E) show that P188 molecules absorb quickly to the air-water interface but to a lesser amount as compared to P188 in “brush”-phase concentration (0.04 mRA/cm). Correspondingly, the surface pressure increases only to π ∼ 8 mN ·m^-1^ and the average layer thickness stabilizes at ∼1.2 nm. However, the lower P188 adsorption apparently allows parallel adsorption of mAb to the interface, which can be deduced from the slow increase of the amide band integral to ∼0.18 mRA/cm. Qualitatively, the changes in layer composition can be confirmed by inspection of the IRRA spectra (Figure 4B): the P188 characteristic ν(C-O-C) vibration at 1100 cm^-1^ decreases, while the deformation vibration of water δ(H-O-H) at 1650 cm^-1^ regresses and the amide bands at 1725 cm^-1^ to 1485 cm^-1^ become apparent.

When the mAb is injected together with P188 in concentration that allow only the formation of a “pancake”-phase (Figure 4C and F), barely any P188 molecule adsorbs to the surface allowing maximal adsorption of the mAb. Thus, the P188 characteristic ν(C-O-C) vibration is only marginally discernible in the spectra, while the amide bands centred at 1550 cm^-1^ and 1650 cm^-1^ are clearly visible (Figure 4C). Figure 4F shows amide band integrals reaching the maximum value of this series of measurement (0.35 mRA/cm) and the integral values in ν(C-O-C) region remain close to zero. Despite the presence of P188 in the solution, mAb is able to adsorb to the air-water interface. This is competitive with slower increase in surface pressure and layer thickness, compared to the cases where the adsorption is dominated by P188. Interestingly, the final average layer thickness is higher (2 nm) than that of a P188 brush (1.5 nm) and that of the mixed P188/mAb layer (1.3 nm).

Notwithstanding the distinct differences in these three measurements, it remains uncertain whether the observed behavior is inherently linked to the phase state of P188 or merely a consequence of concentration dependence. In a pharmaceutical product, based on the three used concentrations, the highly condensed phase of P188 is prevalent.

Results of experiments on the simultaneous injection of mAb and PS20 are shown in the supplementary information (Figure S4). They are very similar to those obtained with “brush”-phase P188, i.e., nearly instantaneous increase in surface pressure, layer thickness and integral of the PS20 marker band and close to zero integral of the amide bands. This shows that also PS20 is able to quickly adsorb to free surfaces and thereby prevent mAb adsorption. PS20 completely shields the surface and mAb adsorption is not possible at any studied PS20 concentration.

#### 2. Antibody injection after formation of a surfactant film

In this series of experiments the surfactants films were formed prior to the mAb injection, i.e. mAb can only adsorb to defects in the layers, would have to replace the surfactants from the interface, or could adsorb to the surfactant layers. These mechanisms can be investigated by stepwise injection of i) the surfactents and ii) the mAb. Because in this experimental desing we can determin starting values in surface pressure, layer thickness and band intensities, we can atribute any changes after mAb injection to film reorganisation.

As in the first set of experiments various P188 concentrations were chosen, such that P188 forms different self-assembled phases at the air-water interface. This allows to investigate whether the poloxamer structure has an impact on mAb interaction. Also, PS20 was tested in various subphase concentrations in Langmuir film balance measurements. The results are shown in the supplementary information (Figure S5). From these experiments it can be deduced that mAb is able to insert into films formed by P188 at concentrations *c*_P188_ < 1 mg/L and films formed by PS 20 concentrations *c*_PS20_ < o.11 mg/L. This shows that PS20 has a higher propensity to shield the interface, i.e., lower amounts of PS20 are necessary to prevent mAb adsorption.

Experiments with selected concentrations were performed with additional IRRAS detection. Figure 5 shows the results of three experiments with representative P188 concentrations, i.e., the time dependent development of surface pressure π, layer thickness *d*, and the integrals of the amide bands as well as the integral of the asymmetrical C-O-C-stretching vibrational band. For related IRRA spectra see Figure S6. The measurements show a similar trend as the previous studies with simultaneous injection: the lower the P188 subphase concentration, the lower is the amount of adsorbed poloxamer and the higher the amount of adsorbed mAb at the air-water interface. P188 existing in the densely packed “brush”-phase (Figure 5A) is able to completely prevent mAb adsorption. Here, neither, the ν(C-O-C) integral nor the surface pressure, nor the layer thickness is influenced by mAb injection. All these measures remain at the values that were adopted by P188 brush phase formation, while the amide band integral is is zero. This shows that mAb is not able to reorganize a or interact with pre-formed P188 brushes at the air-water interface. To test the robustness of the surface shielding effect of a P188 “brush”, the experiment was repeated with ninefold higher mAb concentration (c_sub_ = 45 mg/L). Despite this high mAb concentration in the subphase no mAb was adsorbed to the interface, being blocked by the P188 “brush” phase (Figure S7).

**Figure 5:**
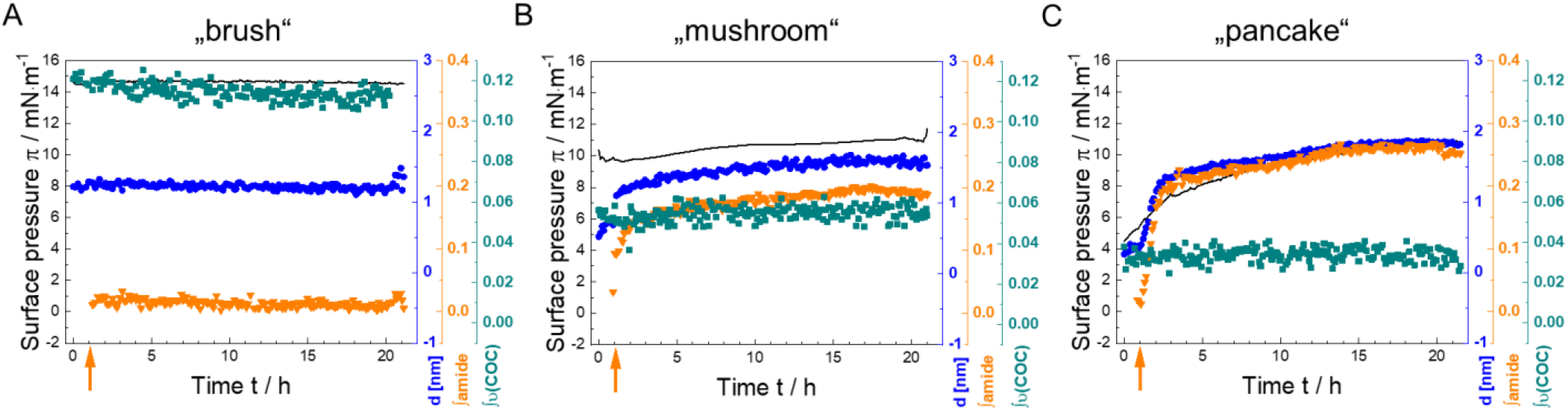
Competitive surface activity experiments of P188 and mAb, using simultaneous infrared reflection-absorption spectroscopy (IRRAS) and surface tension measurements. Different subphase concentrations, (A: “brush” 1 mg/L, B: “mushroom” 0.13 mg/L, C: “pancake” 0.11 mg/L) were used. After one hour of adsorption time, mAb was injected (c_sub_ = 5 mg/L). Measurements were performed on 24.4 mM histidine buffer (pH = 6.0 ± 0.2) at a temperature of T = 20 °C. Surface pressure π (black line), layer thickness *d* 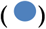, integral of amide-I and amide-II-bands 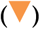 and integral of asymmetrical C-O-C-stretching vibration centred at 1100 cm^-1^ 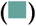 as a function of time (h). The orange-colored arrow shows the injection time of mAb into the subphase.

Nearly the opposite result is found when P188 was allowed to form a more loosely packed “pancake” phase before mAb injection (Figure 5C). In this case significant mAb adsorption takes place despite prior adsorption of P188. Concomitantly, the surface pressure increases from 4 mN/m to 10 mN/m and the layer thickness from 0.3 nm to 1.9 nm. However, mAb does not replace already adsorbed P188 from the interface, as can be deduced from the unaffected and constant integral of the P188 marker band. Consequently, the amount of adsorbed mAb (0.25 mRA/cm) is lower than in the case of simultaneous adsorption of P188 and mAb (0.35 mRA/cm). This shows that even in the pancake phase P188 has a partially protective effect on mAb adsorption.

Similar, but less pronounced effects can be observed when P188 was allowed to form a “mushroom*”* phase before injection of mAb (Figure 5B and Figure S6B). MAb is able to adsorb to the interface despite the presence of P188, while no P188 is displaced from the interface. The concentration of adsorbed mAb (0.2 mRA/cm) is lower than in the case of surface shielding by a “pancake” phase but higher than in the case of simultaneous injection of P188 and mAb (0.18 mRA/cm). Interestingly, the surface pressure is only marginally influenced by mAb adsorption, showing that it is dominated by the more surface active P188.

In addition, we tested the protective effect of a preformed PS20 layer (c_sub_ = 1 mg/L) on mAb adsorption (Figure 6). The results are similar to those obtained for P188 in “brush” phase concentrations. The IRRA spectra in Figure 6A show no amide bands and the respective integrals are roughly zero, showing that no mAb adsorption occurred when PS20 is present at the interface. Consequently, neither layer thickness nor surface pressure are affected by mAb injection into the subphase.

**Figure 6:**
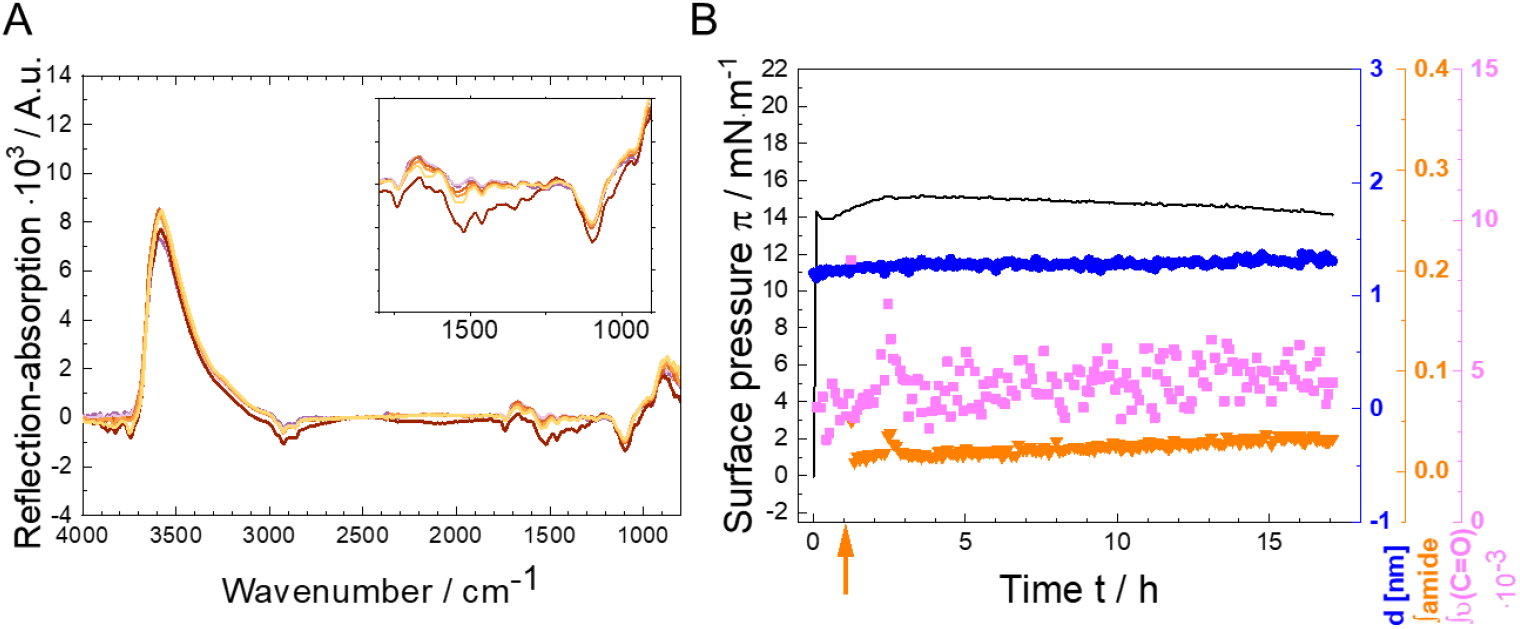
Competitive surface activity experiments of PS20 and mAb, using simultaneous Infrared reflection-absorption spectroscopy (IRRAS) and surface tension measurements. A PS20 subphase concentration of 1 mg/L was used. After about one hour adsorption time mAb was injected (c_sub_ = 5 mg/L). Measurements were performed on 24.4 mM histidine buffer (pH = 6.0 ± 0.2) at a temperature of T = 20 °C. A: IRRA spectra 5 and 30 minutes after PS20 injection and at two to 15.5 hours with additional mAb. The insets shows an enlarged part of the spectra in the wave number range of 1800 cm^-1^ to 900 cm^-1^. The color representation corresponds to different adsorption times. The purple graphs show the spectra of pure PS20. The spectra of PS20 and mAb are shown in orange. Darker shades of purple/orange indicate earlier spectra, while lighter shades of purple/orange indicate later spectra. B: Surface pressure π (black line), layer thickness *d* 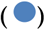, integral of amide-I and amide-II-bands 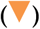 and integral of C=O-stretching vibration centred at 1750 cm^-1^ 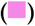 as a function of time (h). The orange-colored arrow shows the injection time of mAb into the subphase while PS20 was injected at *t* = o h.

#### 3. Surfactant injection underneath a preformed antibody layer

To investigate whether an already formed mAb layer can be desorbed from the interface by injection of surfactants, we performed the experiments also in the inversed injection sequence, i.e., mAb was allowed to form an adsorption layer before the surfactants were injected underneath at two different times after film formation has begun. The results are summarized in Figure 7.

**Figure 7:**
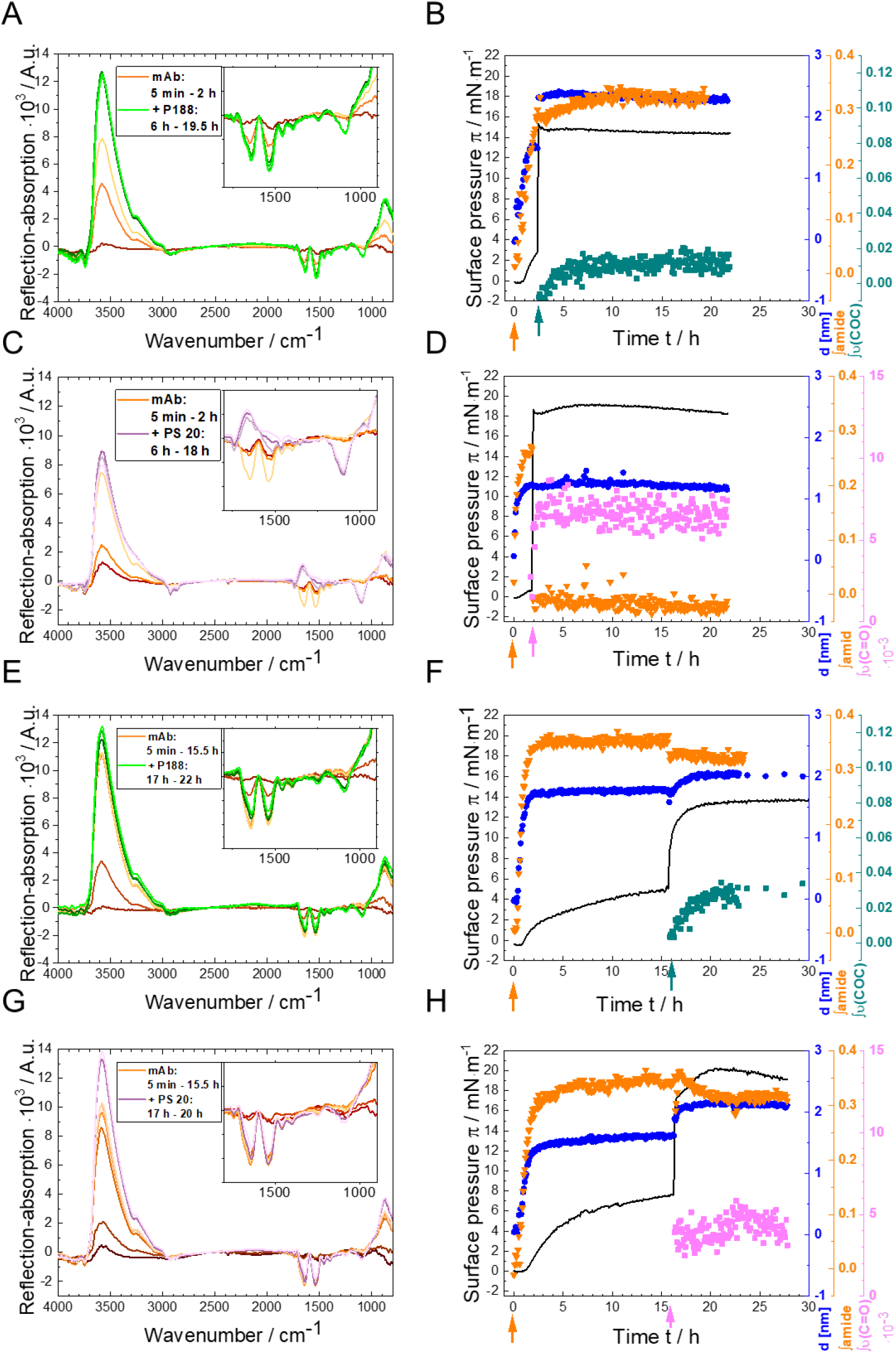
Competitive surface activity experiments of P188 (A,B,E,F), PS20 (C, D,G,H), and mAb, using simultaneous infrared reflection-absorption spectroscopy (IRRAS) and surface tension measurements. A mAb subphase concentration of 5 mg/L was used. mAb was injected at t = 0 h. After about two hours (A-D) or 17 hours (E-H) of mAb film formation the surfactants were injected (*c*_sub_ = 1 mg/L). Measurements were performed on 24.4 mM histidine buffer (pH = 6.0 ± 0.2) at a temperature of T = 20 °C. A, C, E, G: selected IRRA spectra at various times before and after mAb injection. The insets show enlarged part of the spectra in a wavenumber range of 1800 cm^-1^ to 900 cm^-1^. B, D, F, H: Surface pressure π (black line), layer thickness *d* 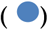, integral of amide I and amide II bands 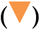 and integral of asymmetrical C-O-C stretching vibration centred at 1100 cm^-1^ 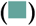 or integral of C=O stretching vibration centred at 1750 cm^-1^ 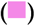 as a function of time. The orange-coloured arrow shows the injection time of mAb into the subphase, the green arrow shows the injection time of P188 into the subphase, the pink arrow shows the injection time of PS20 into the subphase.

In a first experiment P188 was injected two hours after of mAb injection, i.e. immediately after the adsorbed mAb film reaches its equilibrium layer thickness of ∼1.5 nm and the full adsorption of mAb was ensured (Figure 7A,B). P188 was injected in a concentration that would lead to *brush* formation at a pure air-water interface (1 mg/L) and which was shown to inhibit the adsorption of mAb. In the now performed experiment P188 injection leads to an immediate steep increase in surface pressure from 3 to 15 mN/m, a concomitant further increase in layer thickness from 1.5 to 2.3 nm and evolving ν(COC) bands in the IRRA spectra. All these measures indicate that P188 is adsorbed to the interface even though mAb is already present. Interestingly, also the amide band intensity further increases upon P188 injection. This shows that P188 does not replace mAb from the interface, despite its higher surface activity. The rather unexpected increase in amide band intensity might result from secondary structure transitions and/or reorganizations in the already adsorbed protein layer. The amount of adsorbed P188 is low (0.01 mRA/cm). From the integral values it can be concluded that the average P188 surface coverage is only about 10% of that of a P188 in a brush phase (0.11 mRA/cm), even though the surface pressure increases to typical brush phase values (compare to Figure 5A). It can be assumed that P188 forms small patches of brushes that dominate the surface pressure and compresses the adsorbed protein in a manner that influences its conformation. The average layer thickness is, however, dominated by the mAb layer (2.2 nm).

If the same experiment is performed with Ps20 instead of P188 the situation is completely different. The injection of PS20 underneath a pre-formed mAb layer leads to instant desorption of mAb from the interface, as can be concluded from the disappearance of the amide bands from the IRRA spectra (Figure 7C) and the decrease in amide band intensity to zero (Figure 7D), while the C=O stretching vibrational band intensity and the surface pressure increase due to the adsorption of PS20 to the interface. The surface activity of PS20 is about twice as high than that of P188. This may explain that PS20 is able to fully desorb a formed mAb layer from the air-water interface, in contrast to P188. This holds true, when the mAb layer is formed from a 9-fold higher bulk concentration (see Figure S8). Even though surface pressure, interfacial mAb concentration, and the mAb layer thickness were higher in this case, injection of PS20 led to complete desorption of the mAb film from the interface, when injected 2 h after film formation.

These two experiments were repeated with the difference that now the mAb film was allowed to mature for 17 h before the surfactants were injected underneath (Figure 7 E-H). When P188 is injected after 17 h (Figure 7E,F) the film reacts similar to the case were the injection was already carried out after 2 h. That is, an increase in surface pressure to ∼14 mN/m, a slight increase in layer thickness from 1.8 to 2 nm, and the appearance of ν(C-O-C) vibrational bands at 1100 cm^-1^ indicate the adsorption of P188 molecules to the interface, while the mAb is not desorbed. However, the amide band integral decreases slightly from ∼0.35 mRA/cm to ∼ 0.3 mRA/cm upon addition of P188, which one may interpret as a minor part of the mAb being displaced from the surface. Altogether, P188 is not able to fully displace a mAb layer of 17 hours from the air-water interface.

If PS20 is injected after 17 h underneath a matured mAb layer (Figure 7 G,H) the situation changes completely from injection after 2 h. While the surface pressure also increases to ∼20 mN/m, much less PS20 is adsorbed, as can be concluded from the lower integral intensities of the ν(C=O) band and the absence of a ν(COC) vibration in the spectra centred at 1100 cm^-1^. Remarkably, only a minor fraction mAb is displaced from the interface, while the major fraction resists the displacement by PS20. This is similar to the injection of p188 after 17 h but contrary to the injection of PS20 after 2 h, where the protein is completely desorbed from the interface. Apparently, the maturation of the mAb film increases its resistance to desorption from the interface.

With regard to this, it is notable that after injection of mAb the layer thickness as well as the amide band integral reach their equilibrium values after about three hours, while the surface pressure keeps increasing (Figure 7F, H). This might be due to internal film reorganization processes which lead to exposure of hydrophobic moieties of the preotein without recruiting more material. Such reorganization processes have already been described earlier.^48, 49^

The key question concerning the apparent mAb layer maturation is how this layer attains its stability. A slow layer reorganization may include processes like protein unfolding and exposition of more hydrophobic parts to the air-water interface or the formation of intermolecular beta sheets. Similar processes have already been studied by Kanthe *et al*. (2021), who found that unfolding entails exposure of hydrophobic amino acid residues such as isoleucine, leucine and valine, oriented towards the air-side of the surface.^40^ It already has been assumed that monoclonal antibodies exist partially unfolded at the air-water interface and that this denaturation stabilizes the molecule at the surface.^50^ A detailed secondary structure analysis of IRRA amide-bands is beyond the scope of this paper but will be issue of further investigations and may help to explain the maturation process of the mAb layer at the air-water interface.

## SUMMARY AND CONCLUSION

The main results of this study are summarized in Figure 8, which visualized the amount of interfacially adsorbed mAb at different experimental conditions. The conditions where mAb adsorption is prevented or reversed by addition of surfactants can clearly be identified and correspond to the situation where the bars are zero or close to zero. These are: simultaneous or preceding addition of PS20 or P188 in brush forming concentrations and posterior addition of PS20 when the mAb film age is < 2 h. In general, PS20 outperforms P188 in its effectiveness to prevent or reverse mAb adsorption to the air-water interface. Graphically, the different experimental conditions and injection schemes are summarized in Scheme 4.

**Figure 8:**
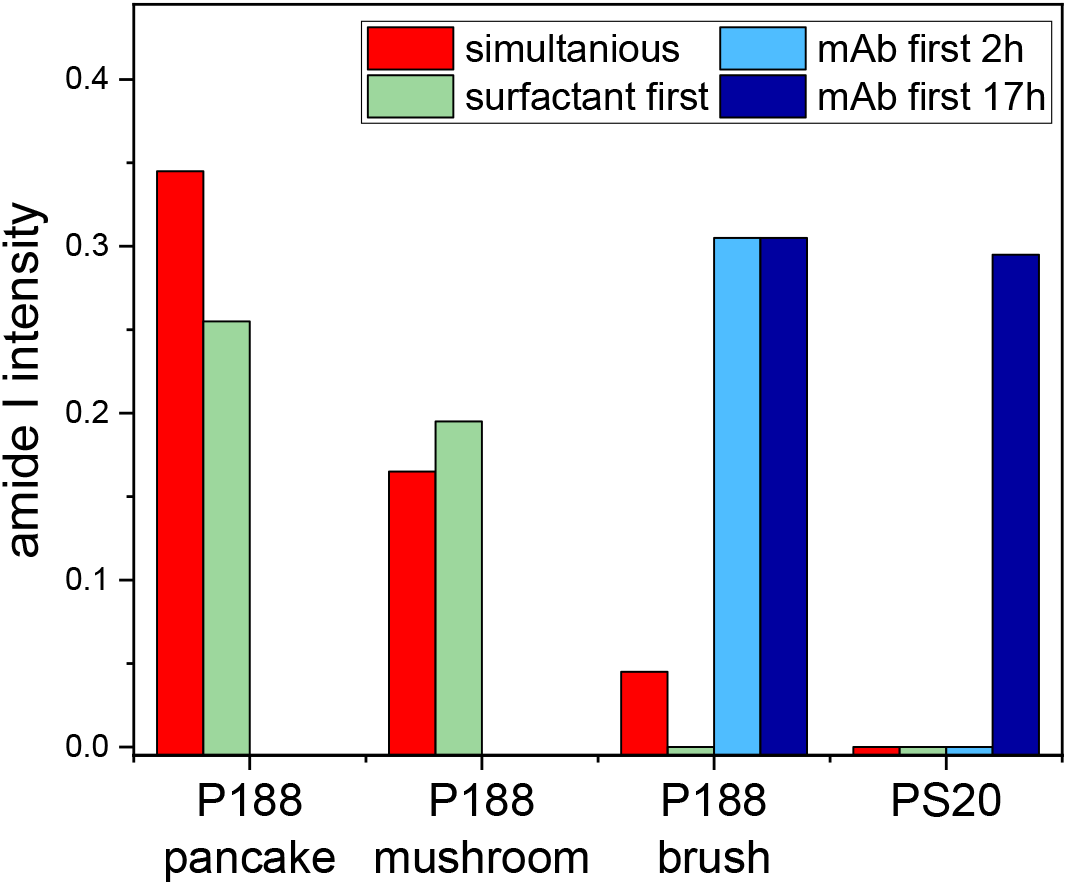
Integral amide band intensity after the competitive adsorption of mAb and P188 or PS20 to the air-water interface. P188 was used in different concentrations (pancake 0.11 mg/L, mushromm 0.13 mg/L, brush 1 mg/L), whereas PS20 was always 1 mg/L. The mAb concentration is always 5 mg/L. The different bar colours denote the applied injection schemes. Red: simultaneous injection of mAb and surfactants, green: injection of mAb underneath a pre-formed surfactant layer, blue: injection of surfactants underneath a pre-formed mAb layer of 2 h (light blue) and of 17 h (dark blue).

In conclusion, adsorption or replacements of the mAb to or from the air-water interface in presence of a surfactant depend on several factors. These are:

i. the nature and amphiphilicity of the chosen surfactant,
ii. the surfactant subphase concentration,
iii. temporal order of film formation, and
iv. the maturation time the mAb film.

The first point is apparent as PS20 is able to shield the surface completely and in addition it is able to fully desorb a previously formed mAb layer up to two hours after its formation. P188 does not show the latter effect in any of our performed experiments and the shielding is incomplete when simultaneous adsorption is allowed. In film balance measurements we furthermore showed that the minimal concentration necessary to prevent mAb adsorption is much lower for PS20 (0.2 mg/L) than for P188 (1 mg/L) (see Figure S5). This effects correlate with the higher surface activity of PS20 over P188. We determined the following order of the surface activities of the tested substances: π_*PS* 20_ > π_*P*188_> π_*mAb*_. This implies that that the surface activity contributes to the effectiveness of the surfactant in replacing the protein from the interface. However, it is not a sufficient criterion to assess whether a substance is replacing another from the interface, since otherwise also P188 should be able to replace mAb.

The surfactant subphase concentration becomes important as it might lead to different phase states and layer structures at the interface, as can be observed for P188. Here, preventing adsorption of mAb to the air-water interface is most effective in the concentration range, where P188 self-assembles in the “brush”-phase. This notably corresponds to the P188 concentrations used in pharmaceutical drugs formulations. The “pancake”-phase of P188 cannot shield the surface and the mushroom phase only partially prevents mAb adsorption. This concentration dependence is represented in Figure 8 by the decrease in red and green bar heights with increasing P188 concentrations.

Furthermore, it could be shown that it is easier to prevent the adsorption of mAb than to reverse it after mAb has once formed an interfacial film. Most effectively, adsorption can be prevented when the surfactant is allowed to occupy the surface before mAb is added to the subphase (green bars in Figure 8). However, simultaneous adsorption of mAb and surfactants leads to similar results (red bars in Figure 8), most likely due to the much faster adsorption kinetics of the surfactants as compared to mAb (Figure 2). Posterior injection leads to mAb desorption only for PS20 and could not be achieved with P188 (blue bars in Figure 8).

For desorption of a once formed mAb layer, the injection time of PS20 is decisive. PS20 injection is necessary during the first two hours after mAb injection to guarantee mAb displacement. A slow reorganization of mAb layer seems to take place at the time scale of several hours, which makes the matured mAb layer more resistant against desorption. Understanding the thermodynamic and structural aspects of these slow reorganization processes in the mAb layer is highly interesting and is currently in the focus of further studies in our group.

**Scheme 4:**
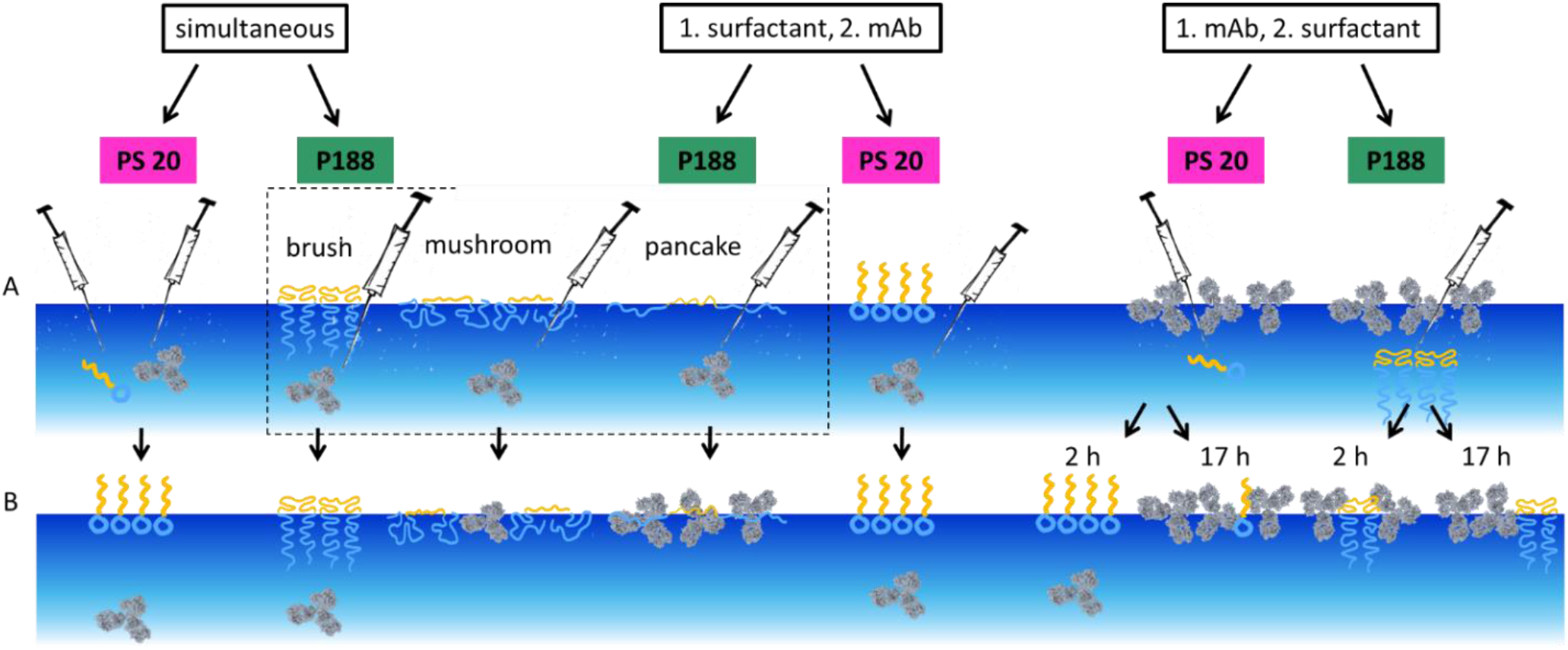
Concluding scheme. A: Injection situation at the beginning of the measurement. B: Results in competitive behaviour. Note: To enhance clarity regarding the phase states of P188, in A only the scheme for injecting mAb beneath a surfactant layer is shown. The schemes for simultaneous injection of mAb and the different P188-phase states are omitted. However, the results in B pertain to both experiments, simultaneous as well as prior P188 injection. For competitive experiments with an existing mAb layer, only the “brush” state of P188 was used. Simultaneous injection of mAb and PS20 or P188 in the “brush” state as well as mAb injection underneath a surfactant layer (PS20 or P188 in the “brush”-phase) leads to a surfactant layer at the air-water interface, mAb remains in the subphase. Simultaneous injection of mAb and P188 in the “mushroom” state as well as mAb injection underneath a P188 “mushroom” state layer leads to a mixed layer of mAb and P188 at the air-water interface. Simultaneous injection of mAb and P188 in the “pancake” state as well as mAb injection underneath a P188 “pancake” state layer leads to a mAb layer at the air-water interface, P188 remains in the subphase. Surfactant injection underneath a mAb layer of two hours leads to a surfactant layer at the air-water interface with mAb remaining in the subphase in case of PS20. In case of P188, a mixed layer of mAb and P188 forms. Similar results are obtained with injection either PS20 or P188 underneath a 17 -hour-old mAb layer.

## ACKNOWLEDGEMENTS

The authors thank Boehringer Ingelheim Pharma GmbH & Co. KG for financial support. EH, CS and DH thank Deutsche Forschungsgemeinschaft (DFG, German Research foundation) - project-ID 436494874 - RTG 2670 ‘Beyond Amphiphilicity (BEAM) for supporting this project.

## SUPPORTING INFORMATION

are available in a separate file: Hingst_mAbadsorption_SI.pdf. They contain additional Figures, calculations of the cmc all used compound, calculation of the area demand of mAb at the interface and images of the instruments.

