## Supporting Information for "Competitive Adsorption of a Monoclonal Antibody and Amphiphilic Polymers to the Air-Water Interface"

Content:

- **Instrument pictures**
- **IRRA band parameters**
- **Area demand of a mAb molecule and surface coverage**
- **Surface activity of pure compound by drop shape tensiometry**
- **cmc calculations**
- **IRRA spectra of pure compounds after subtraction of H2O contribution (to Fig 3, main text)**
- **IRRAS: simultaneous injection of mAb and PS20**
- **Film balance: mAb underneath surfactant film at various surfactant concentrations**
- **IRRA spectra to mAb injection after surfactant film formation (see Fig 5, main text)**
- **Experiments with increase mAb concentration (c = 45 mg/L)**
- **references**

### Instruments pictures

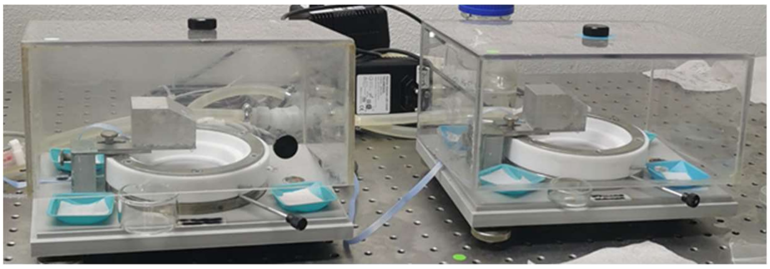

**Scheme S1: Used Langmuir troughs for adsorption measurements.**

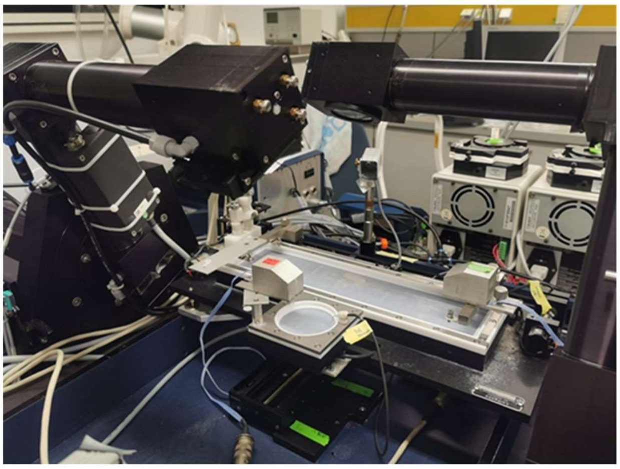

**Scheme S2: Used IRRAS instrument.**

### IRRA band parameters

**Table S1: Integration limits and –methods (OPUS software) for characteristic vibrations of selected functional groups. ν - symmetric stretching vibration, $\nu_a$ - asymmetric stretching vibration, δ - deformation vibration.**

| Substance | Wave-number /cm-1 | Classification | Functional group | Method | Integration limits /cm-1 |
|---|---|---|---|---|---|
| **mAb** | amide I (1658)<br>amide II (1537) | ν (C=O)<br>ν (C-H)<br>δ (N-$R_2$) | amide | B | 1485 - 1725 |
| **P188** | 1100 | $\nu_a$ (C-O-C) | ether | F | 1180 – 1050<br>baseline points:<br>1220; 1180; 1050; 980 |
| **PS20** | 1750 | ν (C=O) | ester | B | 1720 -1760 |

### Area demand of a mAb molecule and surface coverage

For the assessment of the area $A_{molecule}$ that is occupied by a mAb molecule at the air-water interface, the molar volume $V_m$ may be used. According to calculations and experiments of Takashi Imai *et al.* [1], proteins with a molecular weight $M$ of 146'000 g/mol have a molar volume of 102'460 cm³/mol. Based on a maximum layer thickness $d$ of the pure antibody of 1.8 nm and according to the following calculation a value of the available area per molecule is obtained:

$$A_m = \frac{V_m}{d} = \frac{102460\ cm^3/mol}{1.8\ nm} = \frac{1.0246\times10^{26}\ nm^3/mol}{1.8\ nm} = 5.692 \times 10^{25}\ nm^2/mol \qquad \text{(SI Eq.1)}$$

The area per molecule is determined by dividing the result of SI Eq.1 by the Avogadro constant which gives a molecular area of

$A_{molecule}$ = 94 nm² / molecule.

In comparison, the hydrodynamic diameter of mAb can be considered to be about 10 nm [2]. Under simplified assumption of a globular immunoglobulin, this gives a circular cross section area of around 79 nm². When assuming an anisometric mAb (Scheme 1), the occupied surface would be larger. Consequently, a mAb molecule would have an area requirement of at least 79 nm² at an available surface of about 94 nm² per molecule, which fits reasonably well with a literature value of 70 nm² [3]. It can therefore be concluded that the interface is almost entirely covered by the mAb, taking a planar adsorption for granted.

### Surface activity of pure compound by drop shape tensiometry

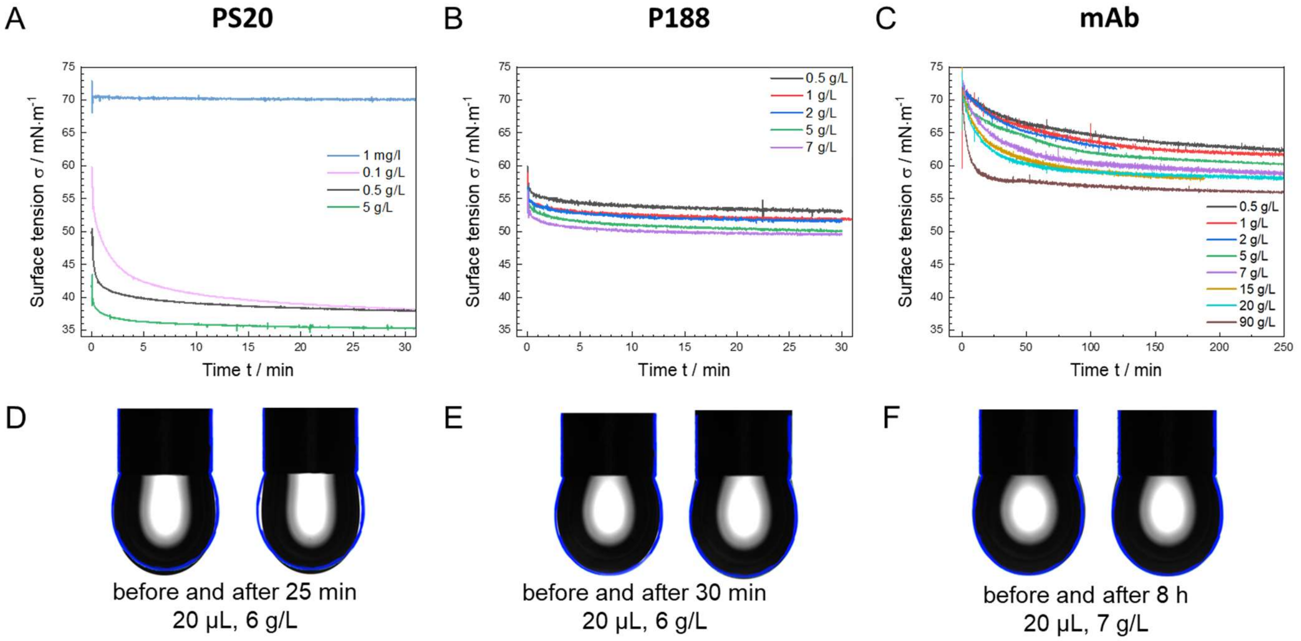


**Figure S1: Surface behaviour of polysorbate 20 (PS20), poloxamer 188 (P188) and monoclonal antibody (mAb) at different subphase concentrations. Measurements were performed on 24.4 mM histidine buffer (pH = 6.0 ± 0.2) at a temperature of T = 20 °C. A-C: Adsorption isotherms on the Drop Shape Tensiometer. D-F: Drop shapes compared to the shape of a water drop (blue outline).**

### cmc calculations

For critical micelle concentration (*cmc*) range determination, the equilibrium surface tension was plotted as a function of the logarithm of concentration and the cmc was determined from the intersection of two linear fits of the data (see SI Fig 2). Literature values are shown on the x-axes [4]. For PS20, a value of 97 µM was experimentally determined, which agrees reasonably well with the literature value of 59 µM [5]. The experimentally determined value for P188 of 3.4 µM is over a factor of 100 lower than the literature value of 480 µM [4,6], which also was the case in prior studies using film balance measurement data [7]. In SI Fig. 2B a kink in the surface tension plots at 53 $mN \cdot m^{-1}$ becomes apparent that has also been found in previous experiments [6]. This behavior might be caused by the phase transition from "brush" to "cigar" which occurs in this region [8].

The concept of a *cmc* is used to describe self-assembly of amphiphilic molecules, in particular surfactants, and therefore does not apply to water soluble proteins like mAbs. Hence it is not surprising that the surface tension values for mAb were not in a state of equilibrium yet, even after 120 minutes, in contrast to the values for both surfactants, which were already taken after 20 minutes.

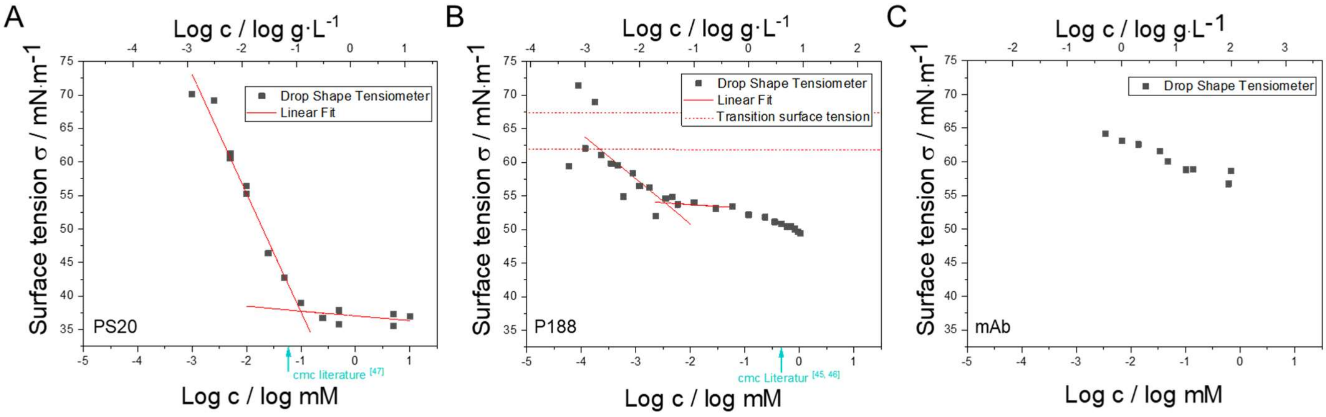


**Figure S2: equilibrium surface tensions as function of concentration of A: polysorbate 20 (PS20) after t = 20 min, B: poloxamer 188 (P188) after t = 20 min, and C: the monoclonal antibody (mAb) solution determined at t = 120 min. Measurements were performed on 24.4 mM histidine buffer (pH = 6.0 ± 0.2) at a temperature of T = 20 °C. Margins of error are within the size of the data points. In A liner fits for determination of the *cmc* are shown. The red dashed lines in B show the transition pressure of the "mushroom"-phase to the "brush"-phase at $\pi$ = 11 $mN \cdot m^{-1}$, which is corresponds to $\sigma$ = 61.8 $mN \cdot m^{-1}$ and of the "pancake" to "mushroom" phase at about $\pi$ = 5 $mN \cdot m^{-1}$, which is corresponds to $\sigma$ = 67.8 $mN \cdot m^{-1}$.**

### IRRA spectra of Fig 3, main text after subtraction of H2O contribution

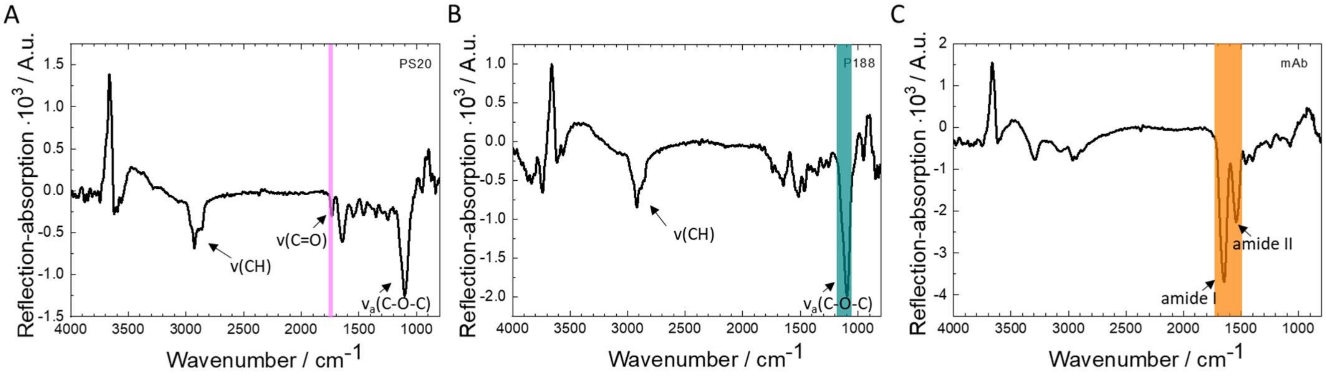


**Figure S3: IRRA spectra of the pure compounds at the air-water interface after subtraction of simulated water spectra as used for further analysis. Measurements were performed on 24.4 mM histidine buffer (pH = 6.0 ± 0.2) at a temperature of T = 20 °C. A, B: after 30 minutes of adsorption, subphase concentration 1 mg/L; C: after 15 hours of adsorption, subphase concentration 5 mg/L. ν, stretching vibration; δ, deformation vibration. Integration limits of the respective bands are shown as a coloured, transparent boxes.**

### IRRAS: simultaneous injection of mAb and PS20

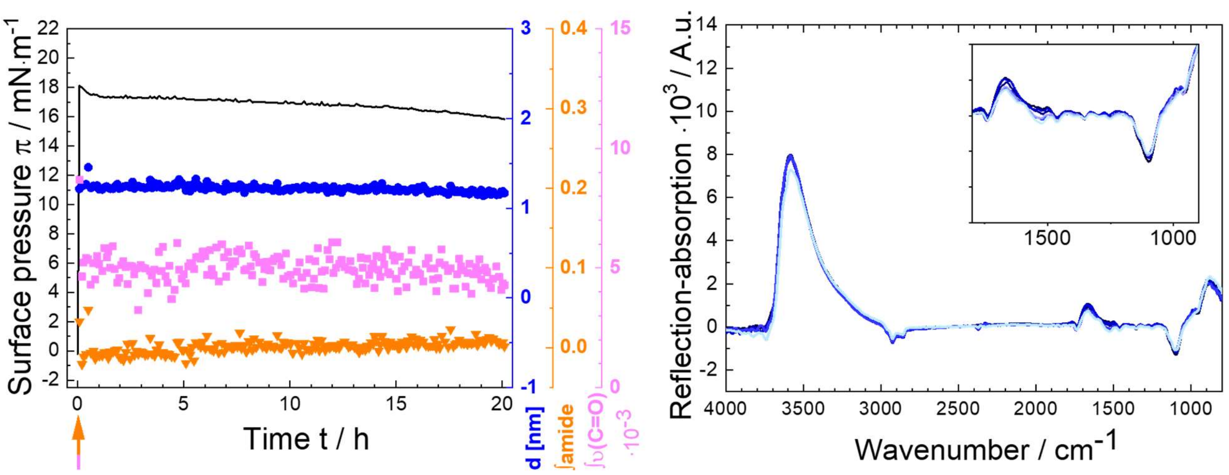


**Figure S4: Competitive infrared reflection-absorption (IRRA) spectroscopic experiments of polysorbate 20 (PS20) and monoclonal antibody (mAb). A simultaneous injection of mAb ($c_{sub}$ = 5 mg/L) and PS20 ($c_{sub}$ = 1 mg/L) was performed. Measurements were performed on 24. 4 mM histidine buffer (pH = 6.0 ± 0.2) at a temperature of T = 20 °C.**
**A: Surface pressure *π* (mN·m$^{-1}$) (black line), layer thickness *d* (●), integral of amide-I and amide-II-bands (▼) and integral of C=O-stretching vibration centred at 1750 cm$^{-1}$ (■) as a function of time / h . The pink-orange arrow indicates the injection time of PS20 and mAb into the subphase. B: IRRA spectra 5 minutes to 15.5 hours after PS20/mAb injection. The inset shows an enlarged part of the spectra of a wave number range of 1800 cm$^{-1}$ to 900 cm$^{-1}$. The color representation corresponds to different adsorption times. Darker shades of blue indicate earlier spectra, while lighter shades of blue indicate later spectra.**

### Film balance: mAb underneath surfactant film at various surfactant concentrations

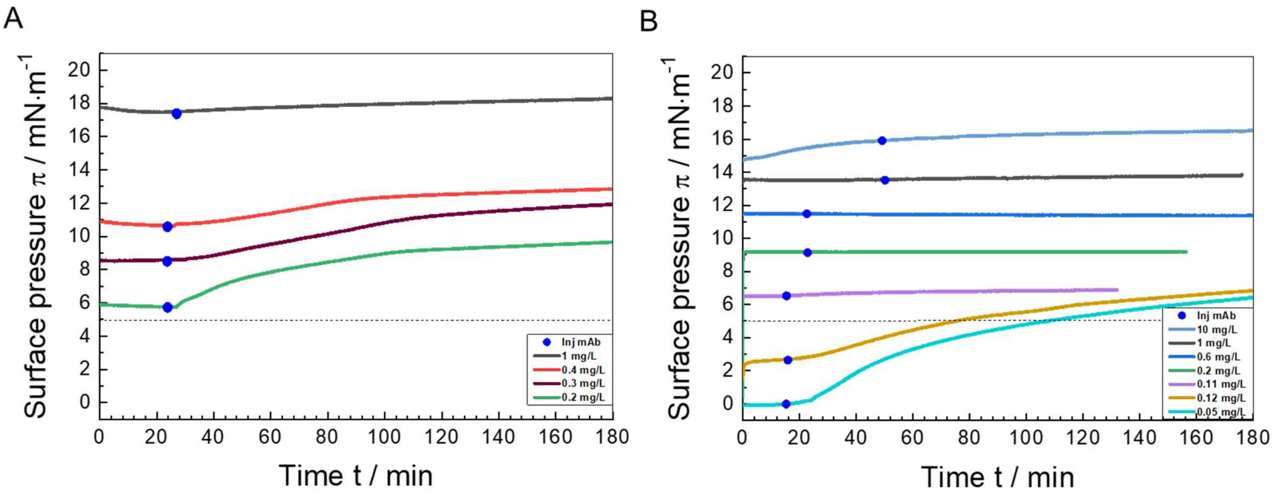


**Figure S5: Competitive adsorption experiments at the Langmuir film balance of (A) poloxamer 188 (P188), (B) polysorbate 20 (PS20) and monoclonal antibody (mAb). Surfactants were injected into the subphase at *t* = 0 h in different concentrations (see legends). Subsequent mAb was injected ($c_{sub}$ = 5 mg/L) at the time indicated by blue filled circles (●). Measurements were performed on 24.4 mM histidine buffer (pH = 6.0 ± 0.2) at a temperature of T = 20 °C. The dotted black line shows the maximal attainable surface pressure of mAb ($c_{sub}$ = 5 mg/L) of about 5 mN·m$^{-1}$.**

### IRRA spectra of mAb injection after surfactant film formation (see Fig 5, main text)

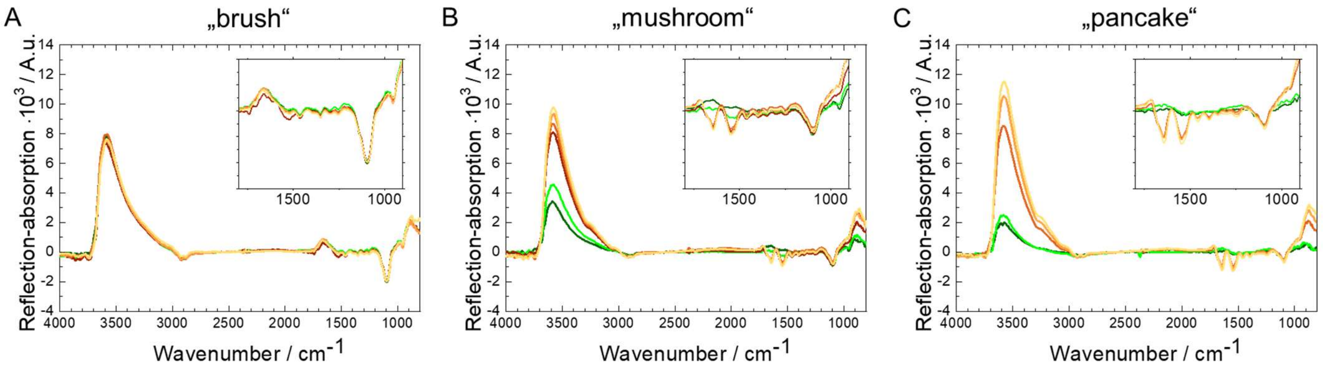


**Figure S6: Competitive infrared reflection-absorption (IRRA) spectroscopic experiments of poloxamer 188 (P188) and monoclonal antibody (mAb). Different subphase concentrations (A: "brush" 1 mg/L, B: "mushroom" 0.13 mg/L, C: "pancake" 0.11 mg/L) were used. After one hour of adsorption time, mAb was injected ($c_{sub}$ = 5 mg/L) followed. Measurements were performed on 24.4 mM histidine buffer (pH = 6.0 ± 0.2) at a temperature of T = 20 °C. A-C: IRRA spectra 5 to 30 minutes after P188 injection and two to 15.5 hours with additional mAb injection. The insets show an enlarged part of the spectra of the wave number range of 1800 cm$^{-1}$ to 900 cm$^{-1}$. The colour representation corresponds to different adsorption times. The green graphs show the spectra of pure P188. The spectra of P188 and mAb are shown in orange. Darker shades of green/orange indicate earlier spectra, while lighter shades of green/orange indicate later spectra.**

### Experiments with increased mAb concentration (c = 45 mg/L)

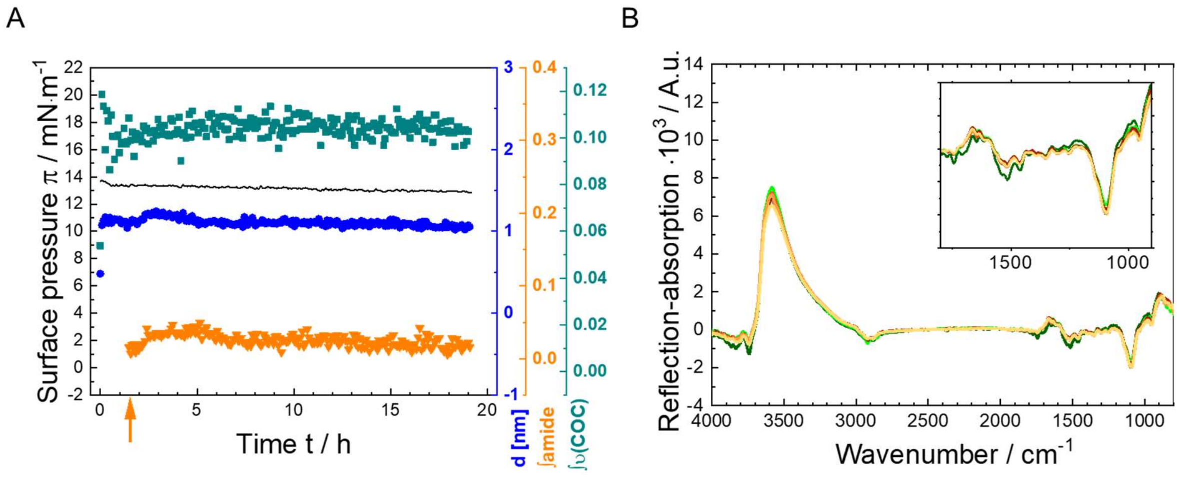


**Figure S7: Competitive infrared reflection-absorption (IRRA) spectroscopic experiments of poloxamer 188 (P188) and monoclonal antibody (mAb). A P188 subphase concentration of 1 mg/L was used leading to formation of a P188 "brush" phase. After about one hour adsorption time mAb was injected ($c_{sub}$ = 45 mg/L). Measurements were performed on 24.4mM histidine buffer (pH = 6.0 ± 0.2) at a temperature of T = 20 °C. A: Surface pressure (black line), layer thickness *d* (●), integral of amide-I and amide-II-bands (▼) and integral of asymmetrical C-O-C-stretching vibration centred at 1100 cm$^{-1}$ (■) as a function of time. P188 was injected at *t* = 0 h into the subphase, the orange-coloured arrow shows the injection time of mAb into the subphase. B: IRRA spectra 5 minutes to 30 minutes after P188 injection and at two to 15.5 hours with additional mAb injection. The inset shows an enlarged part of the spectra in a wave number range of 1800 cm$^{-1}$ to 900 cm$^{-1}$. The colour representation corresponds to different adsorption times. The green graphs show the spectra of pure P188. The spectra of P188 and mAb are**

**shown in orange. Darker shades of green/orange indicate earlier spectra, while lighter shades of green/orange indicate later spectra.**

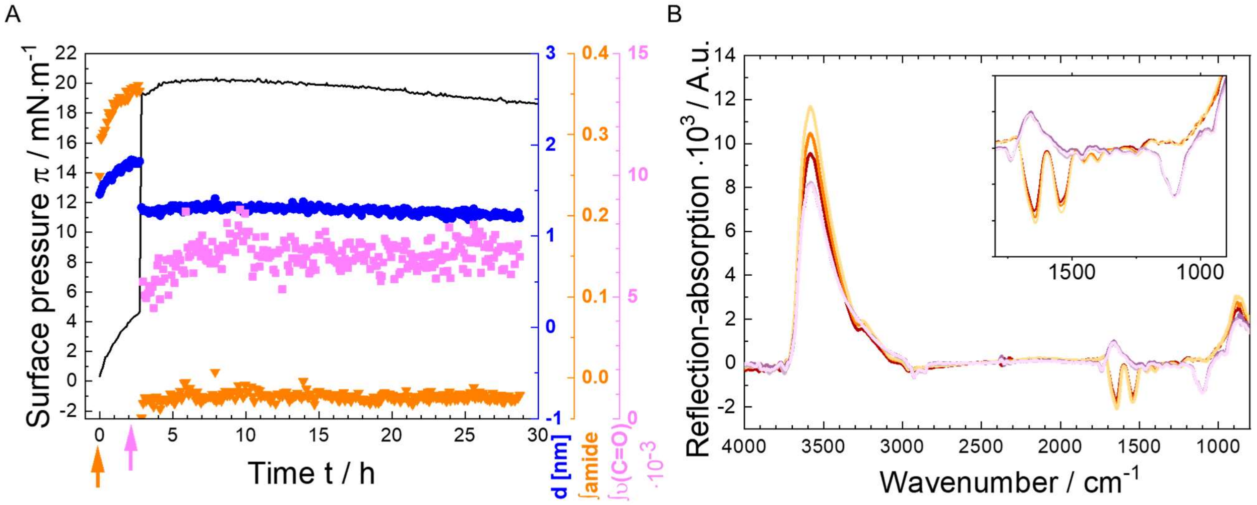


**Figure S8: Competitive Infrared reflection-absorption (IRRA) spectroscopic experiments of polysorbate 20 (PS20) and monoclonal antibody (mAb). A mAb subphase concentration of 45 mg/L was used. After about two hours adsorption time PS20 was injected ($c_{sub}$ = 1 mg/L). Measurements were performed on 24.4 mM histidine buffer (pH = 6.0 ± 0.2) at a temperature of T = 20 °C. A: Surface pressure π (mN·m$^{-1}$) (black line), layer thickness d (●), integral of amide I and amide II bands (▼) and integral of C=O stretching vibration at 1750 cm$^{-1}$ (■) as a function of time (h). The orange-coloured arrow and the pink arrow show the injection time of mAb and PS20 into the subphase, respectively. B: IRRA spectra 5 minutes to two hours after mAb injection and 6 to 15.5 hours with additional PS20 injection. The black box upside right shows an enlarged part of the spectra of a wave number range of 1800 cm$^{-1}$ to 900 cm$^{-1}$. The colour representation corresponds to different time points. The orange graphs show the spectra of pure mAb. The spectra of mAb and PS20 turn to purple. Darker shades of orange/purple indicate earlier spectra, while lighter shades of orange/purple indicate later spectra.**
